# Eukaryotic initiation factor EIF-3.G augments mRNA translation efficiency to regulate neuronal activity

**DOI:** 10.1101/2021.03.15.435499

**Authors:** Stephen M Blazie, Seika Takayanagi-Kiya, Katherine A McCulloch, Yishi Jin

## Abstract

The translation initiation complex eIF3 imparts specialized functions to regulate protein expression. However, understanding of eIF3 activities in neurons remains limited despite widespread dysregulation of eIF3 subunits in neurological disorders. Here, we report a selective role of the *C. elegans* RNA-binding subunit EIF-3.G in shaping the neuronal protein landscape. We identify a missense mutation in the conserved Zinc-Finger (ZF) of EIF-3.G that acts in a gain-of-function manner to dampen neuronal hyperexcitation. Using neuron type-specific seCLIP, we systematically mapped EIF-3.G-mRNA interactions and identified EIF-3.G occupancy on GC-rich 5′UTRs of a select set of mRNAs enriched in activity-dependent functions. We demonstrate that the ZF mutation in EIF-3.G alters translation in a 5′UTR dependent manner. Our study reveals an *in vivo* mechanism for eIF3 in governing neuronal protein levels to control activity states and offers insights into how eIF3 dysregulation contributes to neuronal disorders.

## Introduction

Protein synthesis is principally regulated by variations in the translation initiation mechanism, whereby multiple eukaryotic initiation factors (eIF1 through 6) engage elongation-competent ribosomes with the mRNA open reading frame (Sonenberg and Hinnebusch 2009). eIF3 is the largest translation initiation complex, composed of 13 subunits in metazoans, with versatile functions throughout the general translation initiation pathway (Valasek et al. 2017). Extensive biochemical and structural studies have shown that eIF3 promotes translation initiation by orchestrating effective interactions between the ribosome, target mRNA, and other eIFs (Smith et al. 2016; Cate 2017). Mutations and misexpression of various subunits of eIF3 are associated with human diseases, such as cancers and neurological disorders (Gomes-Duarte et al. 2018), raising the importance to advance mechanistic understanding of eIF3’s function *in vivo*.

Recent work has begun to reveal that different eIF3 subunits can selectively regulate translation in a manner depending on cell type, mRNA targets, and post-translational modification. Interaction of eIF3 RNA-binding subunits with specific 5′UTR stem-loop structures of mRNAs can trigger a translational switch for cell proliferation in human 293T cells (Lee et al. 2015), and can also act as a translational repressor, such as the case for human Ferritin mRNA (Pulos-Holmes et al. 2019). Under cellular stress, such as heat shock, the eIF3 complex circumvents cap-dependent protein translation initiation and recruits ribosomes directly to m6A marks within the 5′UTR of mRNAs encoding stress response proteins (Meyer et al. 2015). Other specialized translation mechanisms appear to involve activities of particular eIF3 subunits that were previously hidden from view. For example, human eIF3d possesses a cryptic mRNA cap-binding function that is activated by phosphorylation and stimulates pre-initiation complex assembly on specific transcripts (Lee et al. 2016; Lamper et al. 2020), while eIF3e specifically regulates metabolic mRNA translation (Shah et al. 2016). These findings hint that many other eIF3 guided mechanisms of cell-specific translational control await discovery.

In the nervous system, emerging evidence suggests that eIF3 subunits may have critical functions. Knockdown of multiple eIF3 subunits impairs expression of dendrite pruning factors in developing sensory neurons of *Drosophila* (Rode et al. 2018). In mouse brain, eIF3h directly interacts with collybistin, a conserved neuronal Rho-GEF protein underlying X-linked intellectual disability with epilepsy (Sertie et al. 2010; Machado et al. 2016). In humans, altered expression of the eIF3 complex in the substantia nigra and frontal cortex correlates with Parkinson’s Disease progression (Garcia-Esparcia et al. 2015). Down-regulation of mRNAs encoding eIF3 subunits is observed in a subset of motor neurons in amyotrophic lateral sclerosis patients (Cox et al. 2010). Furthermore, a single-nucleotide polymorphism located in the intron of human eIF3g elevates its mRNA levels and is associated with narcolepsy (Holm et al. 2015). While these data suggest that eIF3 function in neurons is crucial, mechanistic understanding will require experimental models enabling *in vivo* investigation of how eIF3 affects protein translation with neuron-type specificity.

Protein translation in *C. elegans* employs all conserved translation initiation factors. We have investigated the mechanisms of protein translation in response to neuronal overexcitation using a gain-of-function (*gf*) ion channel that arises from a missense mutation in the pore-lining domain of the acetylcholine receptor subunit ACR-2 (Jospin et al. 2009). The cholinergic motor neurons (ACh-MNs) in the ventral cord of *acr-2*(*gf*) mutants experience constitutive excitatory inputs, which gradually diminish pre-synaptic strength and cause animals to display spontaneous seizure-like convulsions and uncoordinated locomotion (Jospin et al. 2009; Zhou et al. 2017). *acr-2*(*gf*) induces activity-dependent transcriptome changes (McCulloch et al. 2020). However, it is unclear how protein translation conducts the activity-dependent proteome changes that sustain function of these neurons.

Here, we demonstrate that *C. elegans* EIF-3.G/eIF3g regulates the translation efficiency of select mRNAs in ACh-MNs. We characterized a mutation (C130Y) in the zinc-finger of EIF-3.G that suppresses behavioral deficits of *acr-2*(*gf*) without disrupting general protein translation. By systematic profiling of EIF-3.G and mRNA interactions in ACh-MNs, we identified preferential binding of EIF-3.G to long and GC-rich 5’ UTRs of mRNAs, many of which encode modulators of ACh-MN activity. We further provided *in vivo* evidence that EIF-3.G is important for the expression of two mRNA targets dependent on 5′UTRs. Our findings illustrate the selectivity of EIF-3.G in augmenting mRNA translation to mediate neuronal activity changes.

## Results

### A missense mutation in EIF-3.G ameliorates convulsion behaviors caused by cholinergic hyperexcitation

We previously characterized numerous mutations that suppress convulsion and locomotion behaviors of *acr-2*(*gf*) animals (McCulloch et al. 2017). One such suppressor mutation, *ju807*, was found to contain a single nucleotide alteration in *eif-3.G,* encoding subunit G of the EIF-3 complex (Fig. 1A; see Methods). *C. elegans* EIF-3.G is composed of 262 amino acids, sharing overall 35% or 32% sequence identity with human eIF3g and *S. cerevisiae* TIF35 orthologs, respectively (Supplemental Fig. S1A). Both biochemical and structural data show that eIF3g/TIF35 proteins bind eIF3i/TIF34 through a domain in the N-terminus (Fig. 1B) (Valasek et al. 2017). eIF3g/EIF-3.G also has a predicted CCHC zinc finger followed by an RNA recognition motif (RRM) at the C-terminus (Fig. 1B and Supplemental Fig. S1A). The *ju807* mutation changes the second cysteine of the CCHC motif (Cys130, corresponding to Cys160 in human eIF3g) to tyrosine (Fig. 1B). Hereafter, we designate *eif-3.G*(*ju807*) as *eif-3.G*(*C130Y*).

**Figure 1:**
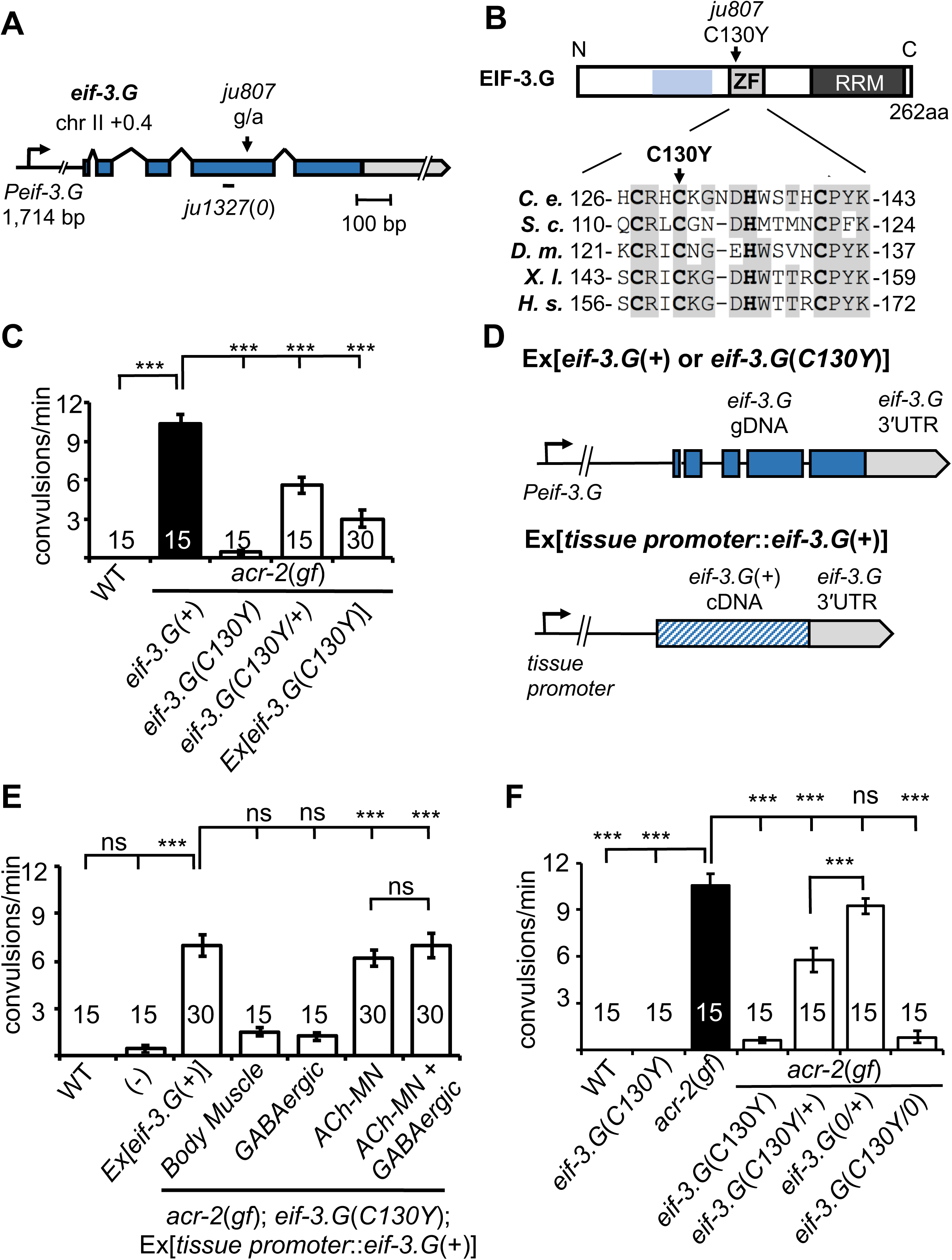
*eif-3.g*(C130Y) suppresses *acr-2*(*gf*) convulsion behavior in the cholinergic motor neurons. **A)** Illustration of the genomic locus of *eif-3.G: Peif-3.G* denotes the promoter, blue boxes are exons for coding sequences and gray for 3′UTR. Arrowhead indicates guanine to adenine change in *ju807;* and short line below represents a 19bp deletion in *ju1327*, designated *eif-3.G*(*0*), that would shift the reading frame at aa109, resulting in a premature stop after addition of 84aa of no known homology. **B)** Illustration of EIF-3.G: shaded blue represents EIF-3.I binding region, ZF for Zinc Finger, RRM for RNA Recognition Motif. Below is a multi-species alignment of the zinc finger domain with bold residues as the CCHC motif and gray for conserved residues. *iu807* causes a C130Y substitution. *C. elegans* (*C. e.*; NP_001263666.1), *S. cerevisiae* (*S. c.*; NP_010717.1), *D. melanogaster* (*D. m.*; NP_570011.1), *X. laevis* (*X. l.;* NP_001087888.1), and *H. sapiens* (*H.s.*; AAC78728.1). **C)** Quantification of convulsion frequencies of animals of indicated genotypes. Ex[*eif-3.G*(*C130Y*)] transgenes (*juEx7015*/*juEx7016*) expressed full-length genomic DNA cloned from *eif-3.g*(*ju807*). **D)** Illustration of *eif-3.G* expression constructs: top shows the transgene expressing genomic *eif-3.G*(+ for wild type and *C130Y* for *ju807*) with the endogenous *eif-3.G* promoter and 3′UTR, and coding exons in blue; bottom shows cell-type expression of *eif-3.G* cDNA driven by tissue-specific promoters (*Pmyo-3*-body muscle, *Punc-25*-GABAergic motor neurons, *Punc-17β* - cholinergic motor neurons). **E)** Quantification of convulsion frequencies shows that convulsion behavior of *eif-3.G*(*C130Y*); *acr-2*(*gf*) double mutants is rescued by transgenes that overexpress *eif-3.G*(+) genomic DNA or an *eif-3.G*(+) cDNA in the ACh-MNs, but not in the GABAergic motor neurons or body muscle. **F)** Quantification of convulsion frequencies in animals of the indicated genotype showing *eif-3.G*(*C130Y*) as a gain-of-function allele. Data in **D-F)** are shown as mean ± SEM and sample size is indicated within or above each bar. Statistics: (***) *P*<0.001, (ns) not significant by one-way ANOVA with Bonferroni’s post hoc test.

Compared to *acr-2*(*gf*) single mutants, *eif-3.G*(*C130Y*); *acr-2*(*gf*) animals exhibited nearly wild type movement and strongly attenuated convulsion behavior (Fig. 1C; Supplemental Videos S1-S3). Additionally, *acr-2(gf)* animals carrying heterozygous e*if-3.G*(*C130Y*/*+*) showed partial suppression of convulsions (Fig. 1C). Overexpression of wild type *eif-3.G* full-length genomic DNA in *eif-3.G*(*C130Y*); *acr-2*(*gf*) double mutants restored convulsions to levels similar to *eif-3.G*(*C130Y*/+); *acr-2*(*gf*) (Fig. 1D-E; Methods). Overexpression of *eif-3.G*(*C130Y*) full-length genomic DNA in *acr-2*(*gf*) single mutants also caused partially suppressed convulsions (Fig. 1C-D). In wild type animals, neither overexpression of *eif-3.G*(+) or *eif-3.G*(*C130Y*) caused any observable effects on locomotion. These data show that *eif-3.G*(*C130Y*) acts in a semi-dominant manner to ameliorate convulsion and uncoordinated locomotion behaviors of *acr-2*(*gf*).

We next determined in which cell types *eif-3.G*(*C130Y*) functions using cell-specific expression analysis (Fig. 1D; also see Methods). *acr-2(gf)* phenotypes arise from a hyperactive ACR-2-containing ion channel expressed in the ventral cord cholinergic motor neurons (ACh-MNs) (Jospin et al. 2009). We found that overexpressing *eif-3.G*(+) cDNA in ACh-MNs (*Punc-17β*) restored convulsions of *eif-3.G*(*C130Y*); *acr-2*(*gf*) animals to a similar degree as those expressing full-length *eif-3.G*(+) under the endogenous promoter (*Peif-3.G*) (Fig. 1E). In contrast, overexpression of *eif-3.G*(+) cDNA in either ventral cord GABAergic neurons (GABA-MNs, *Punc-25*) or body muscle (*Pmyo-3*) in *eif-3.G*(*C130Y*); *acr-2*(*gf*) animals caused no detectable effects (Fig. 1E). Co-expression of *eif-3.G*(+) in both ACh-MNs and GABA-MNs showed similar effects on *eif-3.G*(*C130Y*); *acr-2*(*gf*) animals to that from expressing *eif-3.G*(*+*) in ACh-MNs alone (Fig. 1E). Thus, *eif-3.G*(*C130Y*) functions in ACh-MN to modulate *acr-2*(*gf*) behaviors.

### EIF-3.G(C130Y) selectively affects translation in ACh-MNs

*eif-3.G*(*C130Y*) single mutants exhibit normal development, locomotion, and other behaviors (such as male mating and egg-laying) indistinguishably from wild type animals (Fig. 1F, Supplemental Video S4). Axon morphology and synapse number of ACh-MNs were also normal in *eif-3.G*(*C130Y*) animals (Supplemental Fig. S2A and S2B). To dissect how the C130Y mutation affects EIF-3.G function, we next generated a genetic null mutation (*ju1327*) using CRISPR editing (Fig. 1A and Supplemental Fig. S1A; designated *eif-3.G*(*0*), see Methods). Homozygous *eif-3.G*(*0*) animals arrested development at L1 stage, consistent with EIF-3 complex members being required for *C. elegans* development (Kamath et al. 2003). *eif-3.G*(*0*/+); *acr-2*(*gf*) animals were indistinguishable from *acr-2*(*gf*) single mutants (Fig. 1F). We additionally tested null mutations in EIF-3.E and EIF-3.H, two essential subunits of EIF-3 complex, and found that *acr-2*(*gf*) animals carrying heterozygous null mutations in either *eif-3* subunit gene showed convulsions similar to *eif-3.G*(*0*/+); *acr-2*(*gf*) (Supplemental Fig. S3A). Moreover, hemizygous *eif-3.G*(*C130Y*/*0*) animals are healthy at all stages and suppress behaviors of *acr-2*(*gf*) to levels comparable to *eif-3.G*(*C130Y*) (Fig. 1F). Reducing one copy of *eif-3.H*(*+*) or *eif-3.E*(*+*) in *eif-3.G*(*C130Y*); *acr-2*(*gf*) animals also did not modify the suppression effect of *eif-3.G*(*C130Y*) (Supplemental Fig. S3A). These observations suggest that *eif-3.G*(*C130Y*) retains sufficient function of wild type *eif-3.G*, and likely affects a regulatory activity that is not dependent on EIF-3 subunit dosage.

We considered whether EIF-3.G(C130Y) could alter EIF-3.G protein stability. To test this, we generated single-copy chromosomal integrated transgenes expressing EIF-3.G(WT) or EIF-3G(C130Y) tagged with GFP at the N-terminus under the control of the endogenous *eif-3.G* promoter (Methods and Supplemental Table S1). Both GFP::EIF-3.G(WT) and EIF-3G(C130Y) showed cytoplasmic fluorescence throughout somatic cells and across all developmental stages (Supplemental Fig. S1B). The EIF-3.G(WT)::GFP transgene restored convulsion behavior in the *eif-3.G*(*C130Y*); *acr-2*(*gf*) background and the GFP::EIF-3.G(C130Y) transgene reduced convulsion behavior in the *acr-2*(*gf*) background (Fig. 2A), supporting that GFP::EIF-3.G(WT) and GFP::EIF-3.G(C130Y) retain function. Quantification of GFP levels in the ACh-MNs showed equivalent intensity and localization of GFP::EIF-3.G (WT and C130Y) between wild type and *acr-2*(*gf*) animals (Fig. 2B), indicating that EIF-3.G(C130Y) does not increase EIF-3.G protein stability.

**Figure 2:**
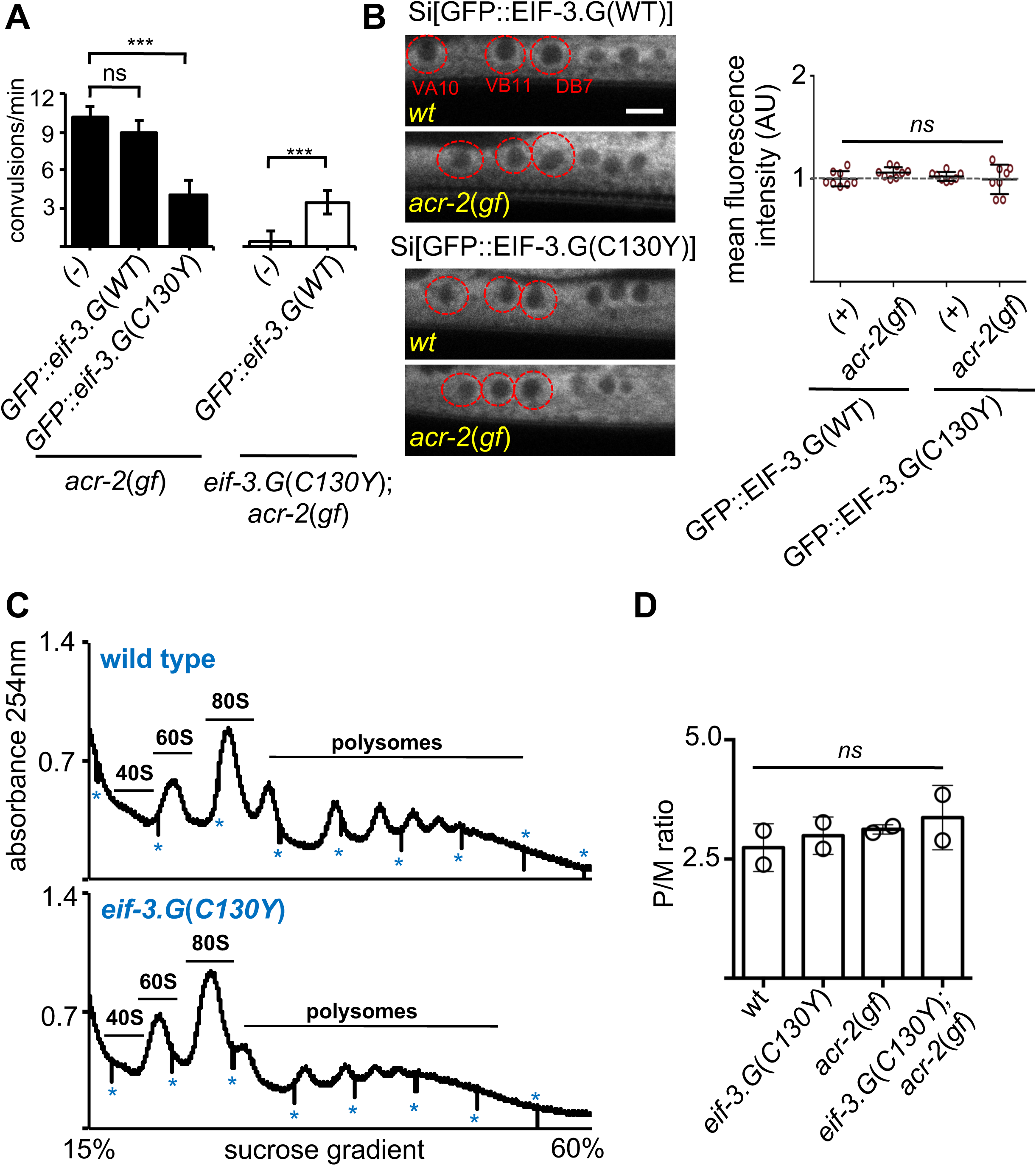
*eif-3.G*(*C130Y*) involves a selective function of EIF-3.G on translational control. **A)** Quantification of convulsion frequency in animals expressing GFP::EIF-3.G(WT) or GFP::EIF-3.G(C130Y) under *Peif-3.G* in the indicated genetic backgrounds. Error bars represent ± SEM with n= 15 per sample. (***) P< 0.001, (ns) not significant, by one-way ANOVA with Bonferroni’s post-hoc test. **B)** EIF-3.G(WT) and EIF-3.G(C130Y) show comparable expression in ACh-MNs. Left are representative single-plane confocal images of EIF-3.G(WT)::GFP or EIF-3.G(C130Y)::GFP driven by the *Peif-3.G* promoter as single-copy transgenes in L4 animals (head to the left). Red circles mark the soma of VA10, VB11, and DB7 ACh-MN, based on co-expressing a *Pacr-2-mcherry* marker. Scale bar = 4 µm. Right: Mean GFP fluorescence intensities (AU) in ACh-MN soma in animals of the indicated genotypes (n= 8). Each data point represents the mean intensity from VA10, VB11, and DB7 neurons in the same animal and normalized to the mean GFP::EIF-3.G intensity in a wildtype background. Error bars represent ± SEM; (ns) not significant by one-way ANOVA with Sidak’s multiple comparisons test. **C)** Representative polysome profile traces from total mRNA-protein extracts of wild type and *eif-3.G*(*C130Y*) single mutant animals. Vertical lines (marked by *) within traces indicate the boundaries of fraction collection. **D)** Polysome::monosome (P/M) ratios calculated based on the area under the respective curves for polysomal and monosome (80S) fractions using two replicates of polysome profiles from total extracts of indicated genotypes. (ns) not significant by one-way ANOVA with Bonferroni’s post-hoc test.

We further assessed whether *eif-3.G*(*C130Y*) alters global translation by performing polysome profile analysis using whole *C. elegans* lysates of L4 stage animals. Both the distribution and ratio of monosomes and polysomes were similar among wild type, *eif-3.G*(*C130Y*), *acr-2*(*gf*) and *eif-3.G*(*C130Y*); *acr-2*(*gf*) animals (Fig. 2C-D), indicating that *eif-3.G*(*C130Y*) possesses normal function in the majority of tissues. It is possible that *eif-3.G*(*C130Y*) suppresses *acr-2*(*gf*) by simply reducing ACR-2 translation. We tested this by examining a functional GFP-tagged ACR-2 single-copy insertion transgene (*oxSi39*). We observed both the levels of ACR-2::GFP fluorescence and post-synaptic localization in ACh-MNs were comparable between wild type and *eif-3.G*(*C130Y*) animals (Supplemental Fig. S3B). These data support the conclusion that *eif-3.G*(*C130Y*) preferentially affects EIF-3’s function in ACh-MNs.

### Regulation of ACh-MN activity by *eif-3.G*(*C130Y*) depends on the mRNA cap-binding protein IFE-1 (eIF4E)

Canonical cap-dependent translation initiation requires the mRNA cap-binding protein eIF4E, which stabilizes a complex that facilitates recruitment of the 40S ribosome to mRNA (Hinnebusch and Lorsch 2012). We tested if the effects of *eif-3.G*(*C130Y*) require *C. elegans* eIF4E homologs. We analyzed convulsion behavior in genetic double and triple mutants of *acr-2*(*gf*) or *acr-2*(*gf*)*; eif-3.G*(*C130Y*) with null mutations in *ife-1*, *ife-2*, or *ife-4* (Fig. 3A; Supplemental Table S1). We detected no effects with *ife-2*(*0*) in any genetic background. *ife-4*(*0*) enhanced convulsions in *acr-2*(*gf*) mutants, but did not alter behavior in *eif-3.G*(*C130Y*); *acr-2*(*gf*) double mutants (Fig. 3A). In contrast, *ife-1*(*0*) significantly restored convulsion behavior in *eif-3.G*(*C130Y*); *acr-2*(*gf*) mutants without influencing the behavior of *acr-2*(*gf*) single mutants (Fig. 3A). Thus, *eif-3.G*(*C130Y*) specifically requires *ife-1*, suggesting cap-dependent translation initiation is likely involved in its modulation of hyperactive ACh-MNs in *acr-2*(*gf*).

**Figure 3:**
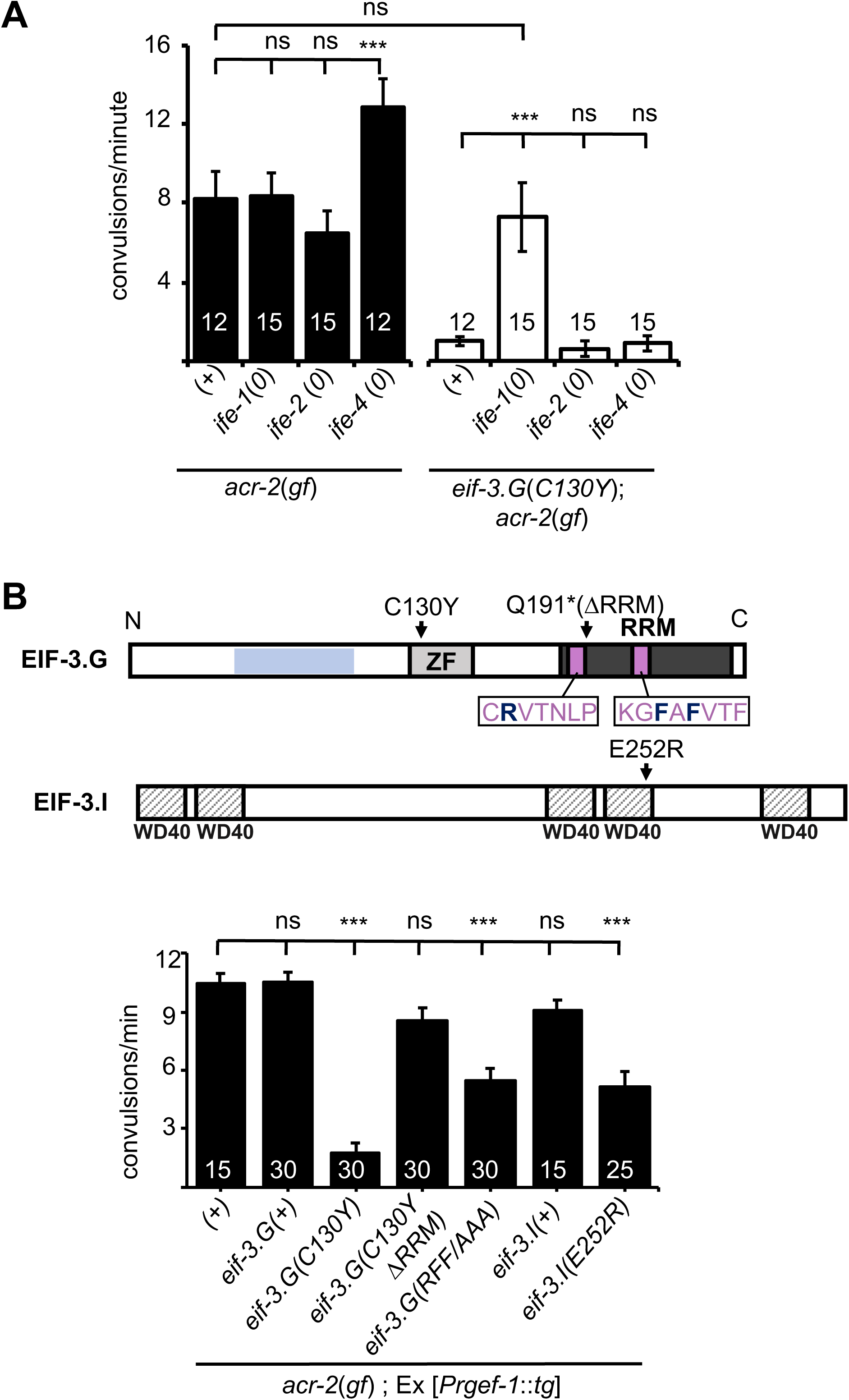
*eif-3.g*(*C130Y*) requires the eIF4E gene *ife-1* and the EIF-3.G RNA-binding domain to suppress *acr-2*(*gf*) behaviors. **A)** Quantification of convulsion frequencies in animals of indicated genotypes. *ife-1, ife-2,* and *ife-4* encode *C. elegans* eIF4E homologs and genetic null mutations in these genes are *bn127*, *ok306*, and *tm684*, respectively (see Supplemental Table S1). **B)** Top illustration of the EIF-3.G protein showing the EIF-3.I binding region (blue), zinc finger (ZF), RRM (dark grey), Q191* mutation in the EIF-3.G(ΔRRM) transgene, RNP motifs (purple), and the RFF residues (bold dark blue) changed to alanine in the *eif-3.G*(*RFF/AAA*) construct. Below is an illustration of *C. elegans* EIF-3.I pointing to the position of E252R within the fourth WD40 domain. Bottom graph is quantification of convulsion frequency in *acr-2*(*gf*) animals expressing *eif-3.G* and *eif-3.I* variants in the nervous system (*Prgef-1*). For **A)** and **B)** Bars represent mean convulsion frequency ± SEM and sample sizes are indicated within or above bars. (***) P< 0.001, (ns) not significant, by one-way ANOVA with Bonferroni’s post-hoc test.

### The activity of EIF-3.G(C130Y) requires its RRM

The RRM located at the C-terminus of eIF3g has been shown to bind RNA in a non-specific manner (Hanachi et al. 1999). To address the role of the RRM in EIF-3.G’s function, we generated a transgene expressing EIF-3.G(ΔRRM) (Fig. 3B; Supplemental Table S1). Expressing EIF-3.G(ΔRRM) under the endogenous promoter *Peif-3.G* in a wild type background did not alter development or locomotion, and also did not rescue *eif-3.G*(*0*) developmental arrest, supporting the essentiality of the EIF-3.G RRM. We then generated a transgene expressing EIF-3.G(C130Y) lacking the RRM domain (C130Y ΔRRM) in neurons of the *acr-2*(*gf*) background (Fig. 3B). In contrast to full-length *eif-3.G*(*C130Y*), *eif-3.G*(*C130Y ΔRRM*) did not alter convulsion behavior of *acr-2*(*gf*) mutants (Fig. 3B), indicating that *eif-3.G*(*C130Y*) function requires its RRM.

Studies on *S. cerevisiae* TIF35/EIF3.G have shown that its RRM promotes scanning of the translation pre-initiation complex through structured 5′UTRs (Cuchalova et al. 2010). Specifically, alanine substitution of three residues in the two ribonucleoprotein (RNP) motifs (K194 in RNP2 and L235 and F237 in RNP1) in TIF35 reduced translation of mRNA reporters carrying 5′UTRs with hairpin structures, without altering the biochemical RNA-binding activity of EIF-3.G/TIF35. Equivalent amino acid residues in *C. elegans* EIF-3.G correspond to R185, F225, F227, which are conserved in human (R242, F282, F284) (Fig. 3B; Supplemental Fig. S1A). To determine whether these residues affect EIF-3.G’s function, we expressed *C. elegans eif-3.G* cDNA with the corresponding amino acids mutated to alanine, designated *eif-3.G*(*RFF/AAA*), in *acr-2(gf)* animals. We detected partial suppression of convulsion behavior in *acr-2*(*gf*) animals (Fig. 3B).

It was also reported that a missense mutation (Q258R) in yeast EIF-3.I/TIF34, located in the sixth WD40 repeat, reduced the rate of pre-initiation complex scanning through 5′UTRs (Cuchalova et al. 2010). To test if *C. elegans eif-3.I* shares similar activities, we made a mutant EIF-3.I(E252R), equivalent to yeast TIF34 (Q258R) (Fig. 3B). In *acr-2*(*gf*) animals, overexpressing *eif-3.I*(*E252R*), but not wild type *eif-3.I*(*+*), caused suppression of convulsions to a similar degree as that by the *eif-3.G*(*RFF/AAA*) transgene (Fig. 3B). These analyses suggest that attenuation of *acr-2*(*gf*) induced neuronal overexcitation may involve regulation of protein translation through modification of 5′UTR scanning rates during translation initiation.

### Both EIF-3.G(WT) and EIF-3.G(C130Y) associate with mRNA 5′UTRs in the cholinergic motor neurons

EIF-3.G may interact with specific mRNAs in the nervous system to regulate cholinergic activity. Therefore, we next searched for mRNAs that are associated with EIF-3.G(WT) and EIF-3.G(C130Y) in the ACh-MNs using single-end enhanced crosslinking and immunoprecipitation (Van Nostrand et al. 2017). We generated single-copy transgenes expressing 3xFLAG-tagged EIF-3.G(WT), EIF-3.G(C130Y), or EIF-3.G(ΔRRM) in the ACh-MNs of *acr-2*(*gf*) animals, with EIF-3.G(ΔRRM) serving to detect indirect crosslinking events. Following cross-linking and immunoprecipitation, we obtained comparable sequencing reads and observed strong correlation between read clusters detected among sets of two biological replicates (Supplemental Fig. S4). We defined EIF-3.G-RNA crosslink sites as clusters of at least 20 high quality reads with at least 1.5 fold change enrichment over the input control (see Methods and Supplemental Table S5). We further defined specific footprints of EIF-3.G(WT) and EIF-3.G(C130Y) by subtracting clusters detected with EIF-3.G(ΔRRM) (Supplemental Table S6, also see Methods). The EIF-3.G specific footprints were primarily located within or near the 5′UTRs of protein-coding genes (5′UTR proximal) (Fig. 4A-B). In total, we detected 231 5′UTR proximal footprints of EIF-3.G(WT) or EIF-3.G(C130Y), which mapped to 225 different genes (Fig. 4C). The number of reads comprising EIF-3.G(WT) or EIF-3.G(C130Y) footprints was similar (e.g. *egl-30*; Fig. 4B) for most of these genes. While some footprints were differentially detected between EIF-3.G(WT) and EIF-3.G(C130Y), this was almost invariably due to small differences in seCLIP signal intensity (read cluster size) between samples close to the 20 read threshold (Fig. 4C), and we therefore did not further pursue its significance. Overall, the footprint map shows that both EIF-3.G(WT) and EIF-3.G(C130Y) predominantly bind within or near the 5′UTRs of 225 genes in the ACh-MNs, hereafter named EIF-3.G targets.

**Figure 4:**
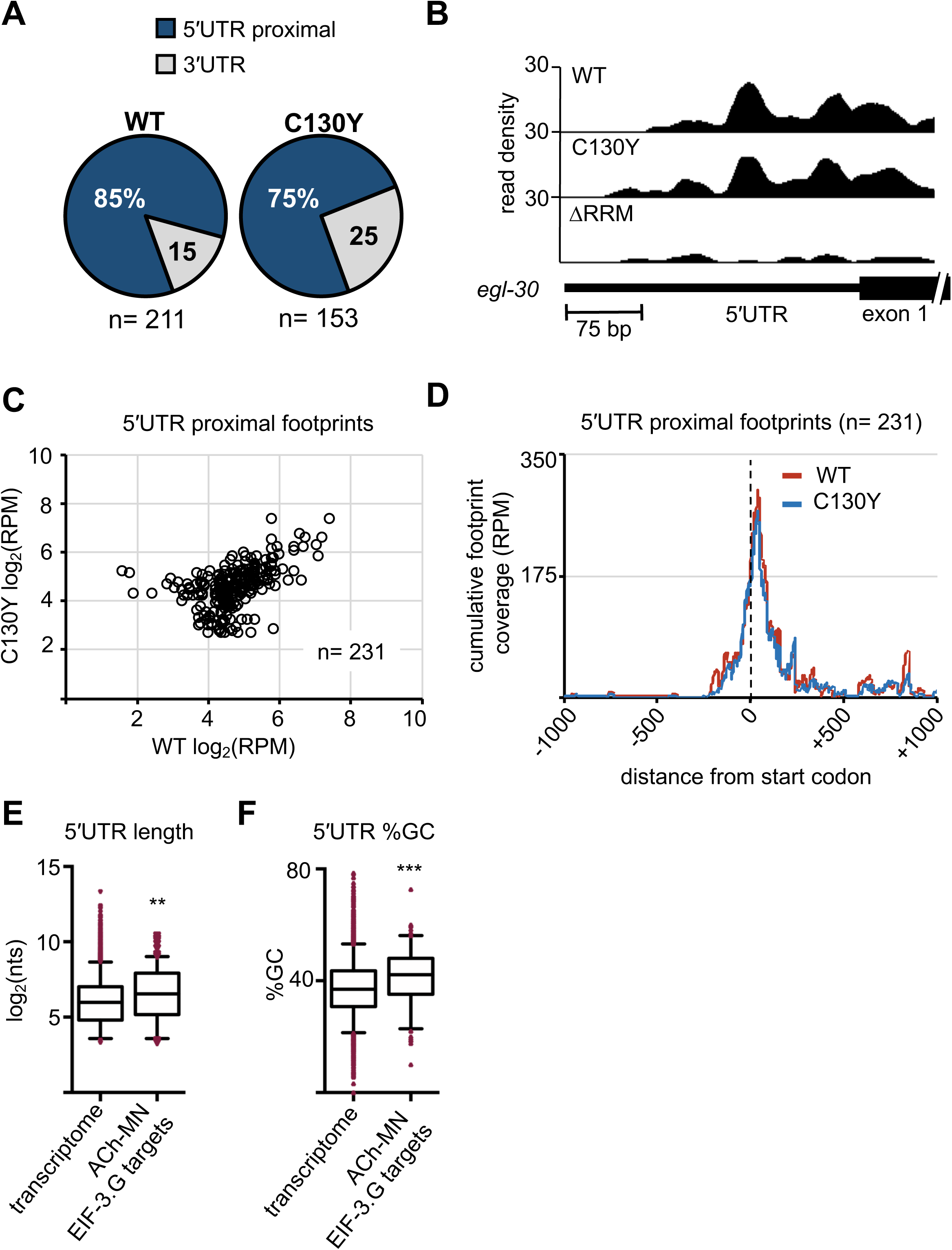
Both EIF-3.G(WT) and EIF-3.G(C130Y) associate with mRNA 5′UTRs in the cholinergic motor neurons. **A)** Pie charts displaying the proportion of EIF-3.G(WT) and EIF-3.G(C130Y) footprints located within each gene feature. **B)** seCLIP read density track of EIF-3.G(WT) and EIF-3.G(C130Y) footprints on the 5′UTR of *egl-30*, compared to the EIF-3.G(ΔRRM) control. **C)** Scatter plot comparing the signal intensity, in reads per million (RPM), of all 231 5′UTR proximal footprints detected in EIF-3.G(WT) or EIF-3.G(C130Y). **D)** Plots show the cumulative coverage of all 5′UTR proximal footprints of EIF-3.G(WT) or EIF-3.G(C130Y) relative to the start codon position. Coverage is presented as reads per million (RPM). **E-F)** Box plots comparing length and GC-content of all 5′UTR sequences of EIF-3.G target mRNAs with annotations (n= 179) to all 5′UTRs in the *C. elegans* transcriptome (n= 10,962). Boxes are 5-95 percentile with outliers aligned in red. Statistics: (***) P< 0.001, (**) P< 0.01 by two-tailed Mann-Whitney test.

In line with a recent report that the human eIF3 complex remains attached to 80S ribosomes in early elongation (Wagner et al. 2020), we observed the bulk of read clusters comprising EIF-3.G(WT) and EIF-3.G(C130Y) footprints mapping between (−)150 to (+)200 nucleotides of the start codon (Fig. 4D). EIF-3.G(WT) and EIF-3.G(C130Y) footprints mapped to similar locations overall (Fig. 4D). Taken together with our finding that *eif-3.G*(*C130Y*) requires its RRM to suppress *acr-2*(*gf*), the seCLIP analysis suggests that the C130Y mutation does not dramatically alter the ability of EIF-3.G to associate with these mRNAs in the ACh-MNs.

### EIF-3.G preferentially interacts with long and GC-rich 5′UTR sequences

5′UTR sequences are widely involved in gene-specific regulation of translation (Pelletier and Sonenberg 1985; Leppek et al. 2018). We next assessed whether the selective role of EIF-3.G in protein translation might correlate with specific sequence features in the mRNA targets expressed in ACh-MNs by examining the length and GC-content of their 5′UTRs. In *C. elegans*, about 70% of mRNAs are known to undergo trans-splicing, and 5′UTRs of mRNAs with trans-splice leaders are usually short, with a median length of 29nt. We compared the EIF-3.G target gene list with a database containing a compilation of *C. elegans* trans-splice events from ENCODE analyses (Allen et al. 2011). We found that 133 of the 225 (59%) EIF-3.G targets are annotated to undergo trans-splicing, which is comparable to that of transcriptome-wide (Allen et al. 2011) (Supplemental Fig. S5A), suggesting that trans-splicing events may not contribute to EIF-3.G’s selectivity on mRNA targets. Interestingly, we found that the trans-spliced 5′UTRs of these 133 transcripts are significantly longer (median length= 43nt), compared with all trans-spliced 5′UTRs in the *C. elegans* transcriptome (median length= 29nt; n= 6,674) (Supplemental Fig. S5B). To assess the GC content for EIF-3.G mRNA targets, we then applied a threshold to the *C. elegans* transcriptome (WS271) annotation defining a 5′UTR as at least 10 nucleotides upstream of ATG, and also selected the longest 5′UTR isoform per gene to avoid redundant analysis of target genes (see Methods). Using this criterion, we identified a 5′UTR for 10,962 different genes in the *C. elegans* transcriptome and for 179 of the 232 EIF-3.G targets in the ACh-MNs. The median 5′UTR among the 179 EIF-3.G target mRNAs was significantly longer (93 nt) and GC-enriched (42%) compared to the *C. elegans* transcriptome median (63 nt and 37% GC; n= 10,962; Figs. 4E-F). We further analyzed the distribution of GC sequences in 5′UTRs, and observed non-random positioning such that some genes were relatively GC-rich near the start codon (e.g. *pmt-2* and *let-607*) and others had enrichment closer to the distal 5’ end (e.g. *unc-43* and *gsa-1*), suggesting that discrete sequence elements in EIF-3.G associated transcripts may regulate translation (Supplemental Fig. S5C).

The incidence of long and GC-enriched 5′UTRs among EIF-3.G associated transcripts led us to speculate a major function of EIF-3.G, in addition to its necessity in general translation initiation, is in the selective regulation of translation. To extend our findings beyond *C. elegans*, we asked if the preferential association of EIF-3.G with these complex 5′UTRs could be conserved in mammals. We analyzed the published eIF3g PAR-CLIP sequencing data from HEK293 cells (Lee et al. 2015) by comparing the 5′UTR lengths of human eIF3g target genes to all genes with 5′UTRs annotated in the hg38 genome. We found that human transcripts associated with eIF3g contained significantly longer and GC-enriched 5′UTRs than average (Supplemental Figs. S5D-E). This analysis lends support for a conserved, specialized role of eIF3g in the translation of transcripts harboring complex 5′UTRs.

### EIF-3.G target mRNAs encode proteins that exhibit activity-dependent expression

To address whether EIF-3.G target mRNAs may preferentially affect specific biological processes, we performed Gene Ontology and KEGG pathway analysis. Significant GO term (Ashburner et al. 2000) enrichment was identified in neuropeptide signaling genes (GO:0050793; 15 genes), which are known to affect *acr-2*(*gf*) behavior (Stawicki et al. 2013; McCulloch et al. 2020), and in stress response genes (GO: 0006950; 28 genes), which could modulate neuronal homeostasis or function under circuit activity changes (Fig. 5A). We also found many EIF-3.G target genes involved in protein translation and protein metabolism processes (GO:0019538; 29 genes; Fig. 5A). Additional enrichment was associated with metabolic components, kinase signaling, and calcium and synaptic signaling pathways (Fig. 5A). Calcium and synaptic signaling genes included the CAMKII *unc-43*, and the G-proteins *egl-30* and *goa-1,* which are all known to regulate ACh-MN synaptic activity (Miller et al. 1999; Richmond 2005; Treinin and Jin 2020).

**Figure 5:**
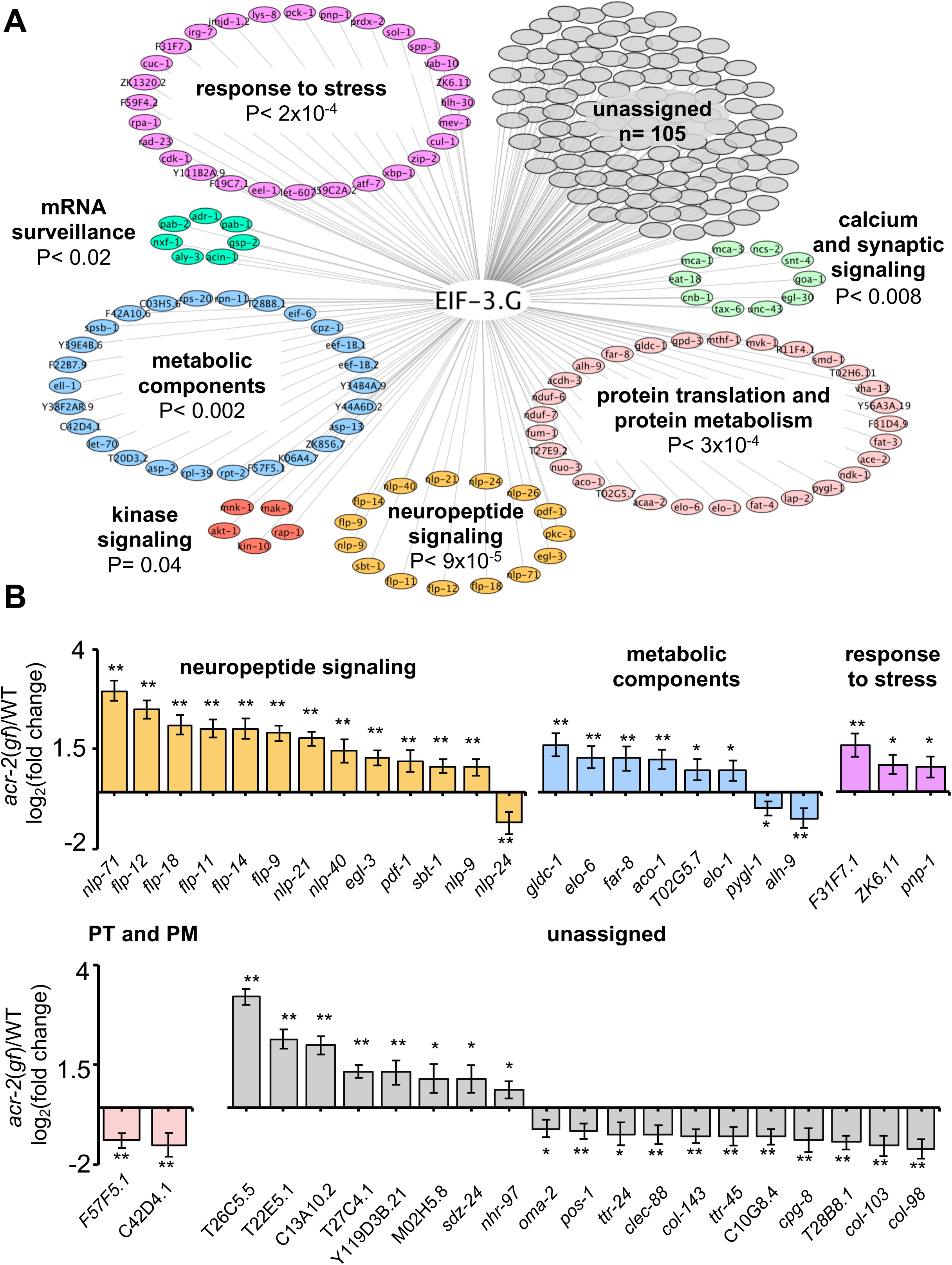
Gene network analyses of EIF-3.G target mRNAs show enrichment in activity-dependent expression. **A)** Cytoscape network of EIF-3.G target genes with enriched GO terms (neuropeptide signaling, response to stress, and protein translation and protein metabolism) or KEGG pathways (calcium and synaptic signaling, metabolic components, MAPK-signaling, and mRNA surveillance). Enrichment P-values are derived from statistical analysis of our EIF-3.G targets (n= 225) in the PANTHER database (Mi et al. 2019). **B)** EIF-3.G target genes exhibiting significant transcript level changes in *acr-2*(*gf*) versus wild type animals as determined from transcriptome sequencing of cholinergic neurons by McCulloch *et al*. PT and PM refers to protein translation and protein metabolism. Differential expression was assessed using DeSeq2 (Love et al. 2014) with significance thresholds of (*) P<0.05 and (**) P<0.01.

To determine if expression of EIF-3.G target mRNAs is regulated in an activity-dependent manner, we next incorporated differential transcript expression data between wild type and *acr-2*(*gf*) from a cholinergic neuron transcriptome dataset (McCulloch et al. 2020). We found that 83% of EIF-3.G target mRNAs in the ACh-MNs are present in the cholinergic neuron transcriptome. Among the 45 genes exhibiting significant expression changes dependent on *acr-2*(*gf*) (Fig. 5B), nearly all neuropeptide signaling transcripts (13 of 15) as well as three stress response genes were upregulated in *acr-2*(*gf*) (Fig. 5B). Genes encoding metabolic components were variably upregulated (e.g. Glycine decarboxylate/*gldc-1*, aconitase/*aco-1*) and downregulated (e.g. glycogen phosphorylase/*pygl-1*, aldehyde dehydrogenase/*alh-9*) (Fig. 5B). These data support the idea that wild type EIF-3.G imparts translational control to activity-dependent expression changes and that EIF-3.G(C130Y) may exert specific regulation to alter their protein expression in ACh-MNs of *acr-2*(*gf*).

### EIF-3.G modulates translation of HLH-30 and NCS-2 in hyperactive ACh-MNs

To experimentally validate that EIF-3.G regulates protein expression from its target mRNAs in the ACh-MNs, we next surveyed a number of candidate genes, chosen mainly based on the availability of transgenic reporters that contain endogenous 5′UTRs (Supplemental Table S1). We identified two genes (*hlh-30* and *ncs-2*) whose expression in ACh-MNs of *acr-2*(*gf*) animals shows dependency on EIF-3.G. *hlh-30* produces multiple mRNA isoforms (Fig. 6A), which encode the *C. elegans* ortholog of the TFEB stress response transcription factor with broad neuroprotective roles (Decressac et al. 2013; Polito et al. 2014; Lin et al. 2018). We observed strong seCLIP signals corresponding to EIF-3.G(WT) and EIF-3.G(C130Y) footprints in the 5′UTR of long isoform d, but not in isoform a (Fig. 6B). The *hlh-30d* mRNA isoform has a 5′UTR of 190nt with 43% GC. Using computational RNA structure prediction (RNAfold), we found that the long *hlh-30d* 5′UTR forms strong stem-loop structures (ΔG= −40.78 kcal/mol) that could affect HLH-30 translation. We examined expression of an HLH-30::EGFP fosmid reporter *wgIs433*, which encompasses the entire *hlh-30* genomic region with *cis*-regulatory elements for all mRNA isoforms (Sarov et al. 2006) (Fig. 6C). HLH-30::GFP was observed throughout the nervous system and primarily localized to cytoplasm in all genetic backgrounds tested. We observed significantly enhanced HLH-30::GFP signals in the ACh-MNs of *acr-2*(*gf*) animals, compared to those in wild type. While *eif-3.G*(*C130Y*) did not alter HLH-30::GFP, it reduced fluorescence intensity in *acr-2*(*gf*) to wild type levels (Fig. 6C). As *hlh-30* transcripts were detected at similar levels in ACh-MNs of wild type and *acr-2*(*gf*) animals (McCulloch et al. 2020), the enhanced HLH-30::GFP signal in *acr-2*(*gf*) likely reflects elevated translation upon neuronal activity changes, which is augmented by EIF-3.G. We also tested a transgenic HLH-30a::GFP reporter expressing *hlh-30a* cDNA driven by the 2 kb sequence upstream of that isoform (Fig. 6D). We found that HLH-30a::GFP intensity was comparable between *acr-2*(*gf*) and *eif-3.G*(*C130Y*); *acr-2*(*gf*) (Fig. 6D). These data strengthen the conclusion that enhanced HLH-30 translation in *acr-2*(*gf*) partly involves the complex 5′UTR of *hlh-30d*.

**Figure 6:**
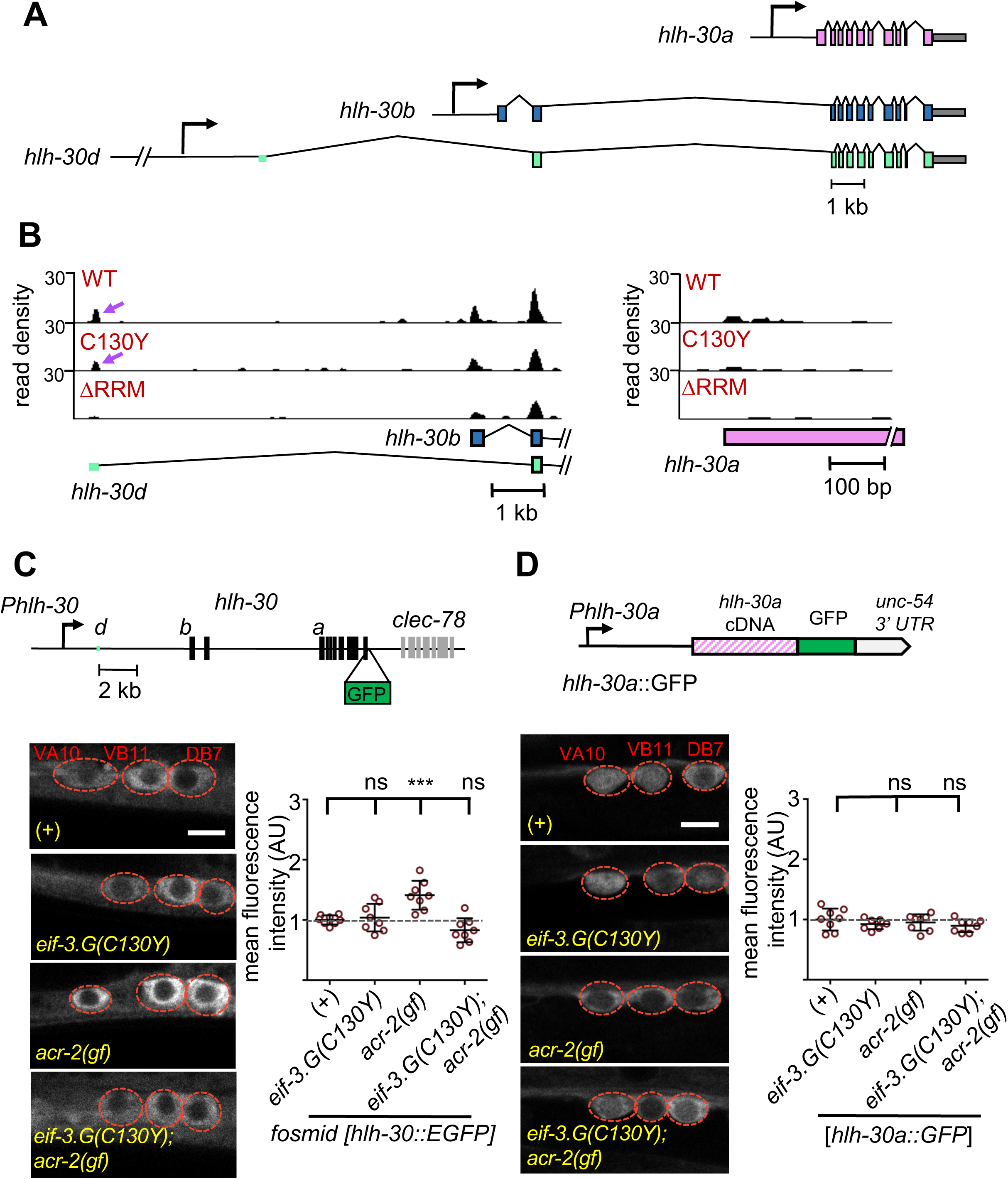
EIF-3.G(C130Y) impairs HLH-30 expression in ACh-MNs of *acr-2*(*gf*) animals. **A)** Gene models of *hlh-30* isoforms *a* (pink), *b* (blue), and *d* (green), with presumptive promoters for each isoform depicted as right-pointing arrows and the 5′UTR of *isoform d* in green to the right of its promoter. **B)** seCLIP read density tracks of footprints on the 5’ end of *hlh-30 isoform b* and *d* (left) and the 5’ end of *hlh-30 isoform a* (right) in each indicated EIF-3.G dataset. Purple arrows show footprints on the 5′UTR of *hlh-30 isoform d*. **C)** Top: Illustration of the *wgIs433* fosmid locus with *hlh-30* coding exons in black and 5′UTR of *isoform d* in green to the right of the promoter. Bottom: Representative single-plane confocal images of the fosmid translational reporter *wgIs433*[*hlh-30*::EGFP::3xFLAG] in ACh-MNs in animals of indicated genotypes. Animals are oriented with anterior to the left. Scale bar = 4 µm. Quantification of GFP intensity (n= 8 for each genotype) is shown on the right. Each data point is the average fluorescence intensity quantified from the three ACh-MN soma per animal and normalized to the mean intensity obtained from *wgIs433* in the wild type background. **D)** Representative single-plane confocal images of *hlh-30a* isoform-specific reporter (*sqIs17*, with the expression construct illustrated above) in ACh-MNs in animals of the indicated genotypes. Scale bar = 4 µm. Average GFP intensities quantified from the soma of ACh-MNs in animals (n= 8) expressing *hlh-30a*::GFP are shown to the right. Data are normalized to the mean *hlh-30a*::GFP fluorescence in a wildtype background. For **C)** and **D)** Red dashes indicate positions of labeled ACh-MN soma. Statistics: (***) P< 0.001, (ns) not significant by one-way Anova with Bonferroni’s post hoc test.

The Neuronal Calcium Sensor protein encoded by *ncs-2* promotes calcium-dependent signaling in ACh-MNs (Zhou et al. 2017). We identified strong and specific association of EIF-3.G(WT) and EIF-3.G(C130Y) overlapping the 5′UTR of *ncs-2* (Fig. 7A). To evaluate NCS-2 expression, we examined a single-copy translational reporter (*juSi260*) expressing NCS-2::GFP under its endogenous promoter (Zhou et al. 2017) (Fig. 7B). NCS-2::GFP localized primarily to the neuronal processes in ventral nerve cord, because of the N-terminal myristoylation motif. Quantification of NCS-2::GFP showed that the fluorescence intensity in *eif-3.G*(*C130Y*); *acr-2*(*gf*) double mutants was significantly reduced, compared to those in wild type, *eif-3.G*(*C130Y*), and *acr-2*(*gf*) (Fig. 7B).

**Figure 7:**
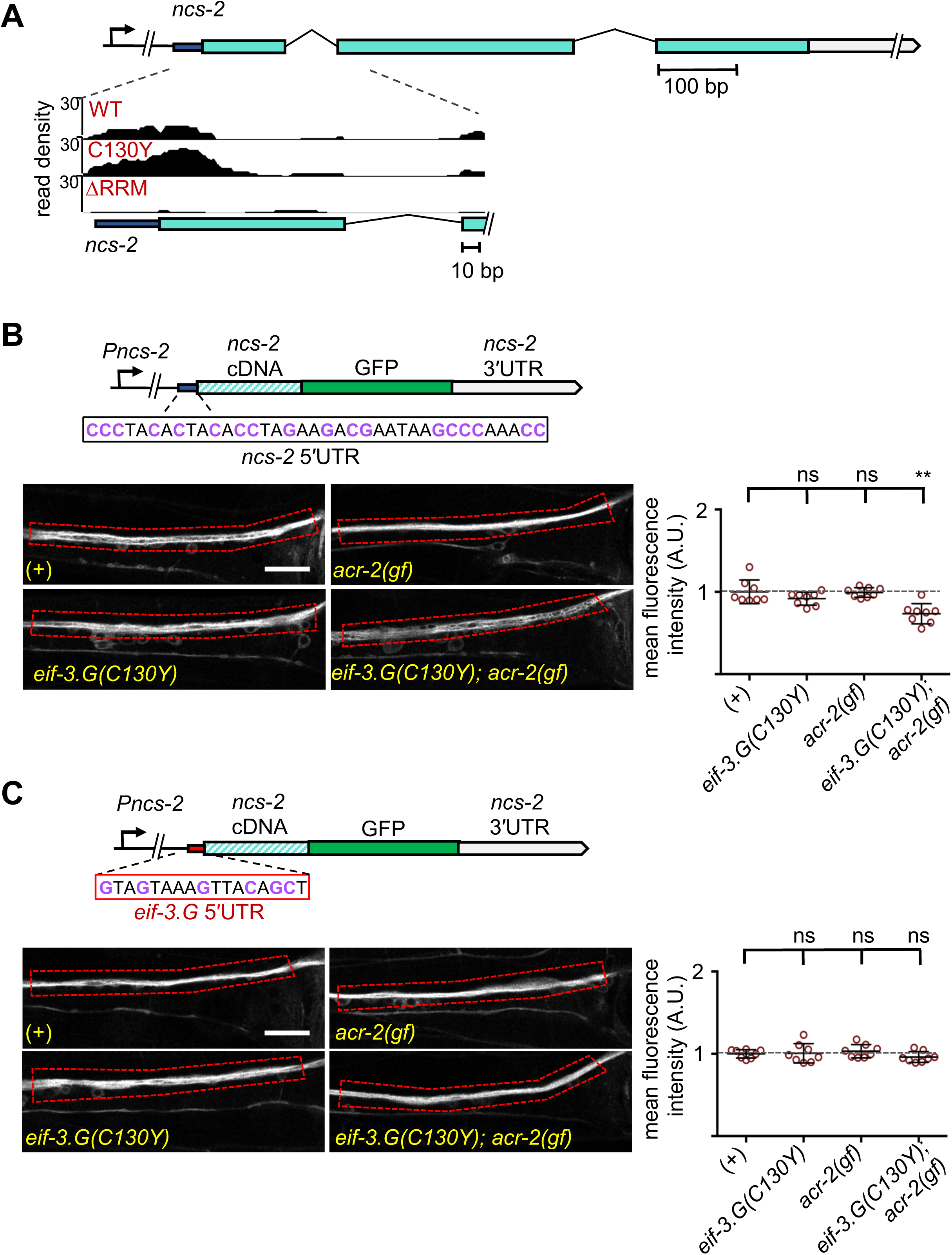
Regulation of NCS-2 expression by EIF-3.G depends on its GC-rich 5′UTR. **A)** Illustration of the *ncs-2* genomic region. Dark blue represents 5′UTR, green boxes are coding exons, and gray is the 3′UTR. The inset below shows the read density track of seCLIP footprints on the 5’ region of *ncs-2* mRNA. **B)** Top: Schematic of the NCS-2(cDNA)::GFP translation reporter, including its 5′UTR (dark blue), driven by the 4kb promoter *Pncs-2*. The 5′UTR sequences are GC rich (purple). *Bottom:* Representative single-plane confocal images of NCS-2::GFP in ventral nerve chord processes in young adult animals of the indicated genotypes. GFP intensity quantification is shown to the right. **C)** Top: The *ncs-2*(5′UTR mutant)::GFP translational reporter has the 5′UTR of *eif-3.G* (red boxed sequence) replacing the *ncs-2* 5′UTR, driven by *Pncs-2*. *Bottom:* Representative single-plane confocal images of ventral nerve chord processes expressing the NCS-2(5′UTR mutant)::GFP translation reporter in young adult animals of the indicated genotypes. GFP intensity quantification is shown to the right. For B) and C) Data points are normalized to the average fluorescence intensity of the respective translation reporter in the wild type background. ROIs used for fluorescence quantification are boxed. Scale bar = 15 µm. Statistics: (**) P< 0.01, (ns) not significant by one-way Anova with Bonferroni’s post hoc test.

*ncs-2* mRNA is SL1 trans-spliced, and the mature 5′UTR has 37 nt that is especially abundant in GC nucleotides (47% GC) (Fig. 7B). Moreover, the *ncs-2* 5′UTR sequence is highly conserved with other nematode species (Supplemental Fig. S6A). By RNAfold prediction we found this sequence could form a strong stem-loop structure (ΔG= −5.10 kcal/mol). To test if NCS-2::GFP expression is regulated specifically through its 5′UTR, we replaced it with the 5′UTR of *eif-3.G*, which is comparatively reduced in GC-content (37% GC) and with much less folding stability (Δ= −1.95 kcal/mol) (Fig. 7C). The *eif-3.G* 5′UTR is also less conserved across nematodes compared to that of *ncs-2* (Supplemental Fig. S6A). We found that the NCS-2::GFP reporter with the 5′UTR of *eif-3.G* was expressed at similar levels in all genetic backgrounds (Fig. 7C).

To further determine the effects of the *ncs-2* 5′UTR in protein translation with neuronal type resolution, we generated a reporter in which the GFP coding sequence was fused in-frame after the first 4 amino acids of NCS-2, which retains the *ncs-2* 5′UTR but disrupts the myristoylation motif, thereby enabling visualization of NCS-2 in ACh-MNs (Supplemental Fig. S6B). Quantification of GFP fluorescence in the cell bodies of VA10, VB11, and DB7 ACh-MN showed significantly reduced expression in *eif-3.G*(*C130Y*); *acr-2*(*gf*) animals (Supplemental Fig. S6B). In contrast, a similar reporter but with the 5′UTR of *eif-3.G* displayed similar GFP levels in all genetic backgrounds (Supplemental Fig. S6C). Therefore, we conclude that *eif-3.G* regulates NCS-2 expression in the ACh-MNs through a mechanism involving its 5′UTR sequence.

## Discussion

The eIF3 complex has been extensively studied for its essential roles in general translation initiation (Cate 2017; Valasek et al. 2017). However, recent work gives support to the idea that eIF3 is also key to many of the specialized translational control mechanisms needed for tissue plasticity *in* vivo (Lee et al. 2015; Shah et al. 2016; Rode et al. 2018; Lamper et al. 2020). Our work expands the landscape of eIF3’s regulatory functions, revealing an *in vivo* role of the eIF3g subunit in stimulating the translation of proteins that mediate neuronal activity changes.

### EIF-3.G ensures the efficient translation of mRNAs with GC-rich 5′UTRs

Our study is the first application of seCLIP-seq to map transcriptome-wide protein binding sites in a specific neuronal subtype (ACh-MNs) in *C. elegans*. With stringent thresholding, we identified 225 genes with strong EIF-3.G occupancy at mRNA 5′ ends. We find that EIF-3.G generally associates with mRNAs harboring long and GC-rich 5′UTRs, implying its RNA-binding function is selective for stimulating translation initiation on 5′ leaders prone to secondary structure or other forms of translation regulation. Our data provide *in vivo* support to the finding that yeast eIF3g/TIF35 promotes scanning through 5′UTRs with stem-loop structures (Cuchalova et al. 2010). The RRM of yeast eIF3g/TIF35 also promotes re-initiation of 40S ribosomes upon terminating at uORF stop codons on GCN4, thereby allowing efficient induction of genes whose translation is regulated by uORFs (Cuchalova et al. 2010). We did not observe uORFs in the 5′UTRs of *ncs-2* or *hlh-30*, suggesting that at least for these mRNAs, *eif-3.G*(*C130Y*) involves reduced scanning through secondary structures or other yet undefined regulatory sequence elements.

It is worth noting that we also found EIF-3.G footprints in 3′UTRs, which could reflect molecular crosstalk between translation initiation and 3′UTR factors, given their proximity in the closed loop translation model (Imataka et al. 1998; Wells et al. 1998). EIF-3.G might anchor the closed-loop mRNA form that stimulates multiple rounds of translation, as was shown to be the case with eIF3h (Choe et al. 2018). It is also possible that EIF-3.G cooperates with 3′UTR interacting factors that regulate gene expression, as several *C. elegans* translation initiation factors co-immunoprecipitated with the miRISC complex (Zhang et al. 2007) and accumulating evidence supports interplay between various translation factors and RISC proteins that mediate translational repression by microRNAs (Ricci et al. 2013; Fukaya et al. 2014; Gu et al. 2014). Thus, further analysis is needed to examine the biological meaning of EIF-3.G association with 3′UTRs.

### The EIF-3.G zinc finger conveys a selective function to translation initiation

The function of the zinc finger of eIF3g remains undefined. Through analysis of EIF-3.G(C130Y), our data provides *in vivo* insights that the zinc finger contributes to translation efficiency of mRNAs harboring complex 5′UTRs. We establish that EIF-3.G(C130Y) behaves as a genetic gain-of-function mutation without disrupting EIF-3 assembly or otherwise impairing general translation, measured by both polysome levels and the health of cells, tissues and animals. The effect of EIF-3.G(C130Y) on *acr-2*(*gf*) behaviors depends on the EIF-3.G RRM as well as the cap-binding factor IFE-1/eIF4E, suggesting that assembly of the pre-initiation complex on mRNA is required for EIF-3.G(C130Y) function. While we did not observe significant mis-positioning of EIF-3.G-mRNA interactions by EIF-3.G(C130Y), we acknowledge that seCLIP may not have the resolution required to reveal subtle differences in crosslinking sites caused by the C130Y alteration. Together, our data is consistent with a model where EIF-3.G(C130Y) imposes a translational stall after EIF-3 complex assembly and mRNA recruitment. In this view, we speculate that the Zinc Finger domain of EIF-3.G mediates interactions with other proteins, such as the ribosome, that critically regulate translation events after mRNA binding. In support of this model, yeast eIF3g/TIF35 was found to directly bind to small ribosomal protein RPS-3, though the molecular basis for mediating this interaction is not identified (Cuchalova et al. 2010). Further studies are required to address the precise molecular mechanism by which the EIF-3.G zinc finger imparts regulatory control over translation initiation.

### EIF-3.G targets the translation of mRNAs that modulate neuronal function

Our study was driven by the genetic evidence that *eif-3.G*(*C130Y*) ameliorates convulsion behavior caused by the hyperactive ion channel ACR-2(GF). We show that EIF-3.G(C130Y) retains essential EIF-3.G function, yet it augments protein translation on select mRNAs in ACh-MNs, as evidenced by its effects on NCS-2 and HLH-30 expression. We previously reported that complete loss-of-function of *ncs-2* strongly suppresses *acr-2*(*gf*) behaviors to a similar degree as *eif-3.g*(*C130Y*) (Zhou et al. 2017). However, 50% reduction of *ncs-2* expression does not cause detectable consequences and complete loss-of-function in *hlh-30* also has no effects in either wild type or *acr-2(gf)*. Thus, the small reduction of NCS-2 and HLH-30 waged by *eif-3.G*(*C130Y*) is unlikely to account for the full extent of phenotypic suppression of *acr-2*(*gf*). Our seCLIP data also revealed EIF-3.G interactions with many other genes that differentially impact *acr-2*(*gf*) behavior (e.g. neuropeptide *flp-18*, endopeptidase *egl-3*) and cholinergic activity (e.g. G proteins *goa-1*, *egl-30*). Interestingly, many of the pre-synaptic genes that regulate *acr-2*(*gf*) behavior, such as *unc-13*/Munc13, *unc-17/VAChT* (Zhou et al. 2013; Takayanagi-Kiya et al. 2016; McCulloch et al. 2017), do not have EIF-3.G footprints. Thus, our data is consistent with a model where *eif-3.G*(*C130Y*) ameliorates behaviors of *acr-2*(*gf*) through the cumulative changes of select ACh-MN activity regulators.

### *eif-3.G* function may be specialized for activity-dependent gene expression

The eIF3 complex is widely implicated in brain disorders, and deregulated eIF3g is specifically linked to narcolepsy (Gomes-Duarte et al. 2018). However, given the essential role of eIF3 in protein translation in all tissues, investigation of its functions in the nervous system remains limited. Our results reveal that EIF-3.G permits normal activity-dependent protein expression changes, and suggest that dysregulated EIF-3.G might potentiate aberrant neuronal behavior in disorders such as epilepsy by altering the neuronal protein landscape. It is worth noting that pore-lining mutations in human nicotinic receptors that occur at similar positions as *acr-2*(*gf*) are causally linked to epilepsy (Xu et al. 2011). We speculate that EIF-3.G may be a potential target for intervention of disorders involving abnormal neurological activity.

In summary, our findings echo the general notion that fine-tuning the activity of essential cellular machinery, such as ribosomes and translation complexes holds the key to balance cellular proteome under dynamic environmental challenges or disease conditions. Emerging studies from cell lines show that stress conditions can induce post-translational modification of eIF3 subunits (Lamper et al., 2020) or cap-independent interactions with mRNAs to modify proteomes (Meyer et al. 2015). Through characterization of the G subunit of eIF3, we reveal the first mechanistic insights into how the eIF3 complex regulates neuronal activity. It is likely that individual eIF3 subunits could each possess unique functions relevant in certain contexts, altogether providing the eIF3 complex with extensive utility to remodel the proteome in response to changing cellular environments.

## Methods

### *C. elegans* genetics

All *C. elegans* strains were maintained at 20°C on nematode growth media (NGM) plates seeded with OP50 bacteria (Brenner 1974). Compound mutants were generated using standard *C. elegans* genetic procedures and strain genotypes are listed in Supplemental Table S1. Primers for genotyping are in Supplemental Table S2.

### Identification of *eif-3.g(ju807)*

We employed a custom workflow on the GALAXY platform to identify SNPs unique to strains containing suppressor mutations of *acr-2*(*gf*), compared to the N2 reference strain (McCulloch et al. 2017). Following SNP mapping using genetic recombinants, we located *ju807* to *eif-3.G* on chromosome II. We then performed transgenic expression experiments and found that both over-expression and single-copy expression of *eif-3.G*(+) in *ju807; acr-2(gf)* animals restored convulsions.

### Quantification of convulsion behavior

Convulsions were defined as contractions that briefly shorten animal body length, as previously reported (Jospin et al. 2009)(Supplemental Video S2). L4 larvae were cultured overnight on fresh NGM plates seeded with OP50 bacteria at 20°C. The following day, each young adult was moved to a fresh seeded plate, and after climatized for 90 seconds, convulsions were counted over a subsequent 90 seconds. The average convulsion frequency represented data over 60 seconds. All statistical tests were performed using GraphPad Prism6 software and P-values <0.05 were considered significant.

### CRISPR-mediated genome editing

We used a previously described method (Dickinson et al. 2013) with minor modifications to generate *eif-3.G(ju1327)* deletion allele. Briefly, we designed sgRNA target sequence CAATTCACAAGAAATCGCGC, and cloned it into a Cas9-sgRNA expression construct pSK136 (derived from pDD162, with site-directed mutagenesis). A DNA mixture containing 50 ng/µl pSK136, 1ng/µl *Pmyo-2::mCherry* (pCFJ90), and 50 ng/µl 100bp ladder (Invitrogen, Carlsbad, CA) was microinjected into N2 adults. We screened F2 progenies from F1 animals carrying the co-injection marker for deletions in *eif-3.G* and identified a 19 bp deletion, designated *ju1327*. Heterozygous *ju1327* was twice outcrossed to N2 and then crossed to the *mnC1* balancer for stable strain maintenance (CZ22974).

### Molecular biology and transgenesis

All transgene constructs were cloned using the Gateway™ cloning system (Invitrogen, Carlsbad, CA) or Gibson Assembly^®^ (NEB, Ipswich, MA), unless otherwise noted. Primers used in their construction are detailed in Supplemental Table S3. For single-copy insertion transgenes, we used a previously described CRISPR/Cas9 method to integrate a single genomic copy on chromosome IV (Andrusiak et al. 2019). For extrachromosomal transgenes, we microinjected a DNA mixture containing 2 ng/µl transgene plasmid, 2.5 ng/µl pCFJ90(*Pmyo-2*::mCherry), and 50 ng/µl 100 bp ladder (Invitrogen, Carlsbad, CA) into young adults, following standard procedure (Mello et al. 1991).

To generate the *eif-3.G(+)* or *eif-3.G*(*C130Y*) genomic constructs (pCZGY3006 or pCZGY3007), we amplified a 2,223 bp region from genomic DNA of N2 or CZ21759 *eif-3.G*(*C130Y*); *acr-2*(*gf*), respectively, which includes 1,714 bp upstream of the start codon of isoform A (F22B5.2a.1) and 331 bp downstream of the stop codon, and cloned the amplicon into the PCR8 vector (Invitrogen, CA).

To generate all *eif-3.g* cDNA expression clones, we made mixed-stage cDNA libraries with poly-dT primer for N2 or CZ21759 using Superscript III (ThermoFisher Scientific, San Diego, CA). We then amplified and *eif-3.G* cDNA using primers for the SL1 trans-splice leader (YJ74) and *eif-3.G* isoform A 3′UTR (YJ11560) and Phusion polymerase (Thermo Fisher Scientific, San Diego, CA). The cDNA clones in PCR8 vector were then used to generate tissue-specific expression constructs using Gateway™ cloning destination vectors (pCZGY1091 for *Punc-17β*, pCZGY925 for *Pmyo-3*, pCZGY66 for *Prgef-1*, and pCZGY80 for *Punc-25*).

We used PCR site-directed mutagenesis, in which the nucleotide changes are introduced by the primers to generate the *Prgef-1*::*eif-3.G*(ΔRRM) and *Prgef-1*::*eif-3.G*(C130Y ΔRRM) constructs (pCZGY3026 and pCZGY3027, respectively) with primers YJ11561 and YJ11562 on the templates pCZGY2715 and pCZGY2716, respectively. The *Prgef-1*::*eif-3.G*(RFF/AAA) construct (pCZGY3512) was generated by two rounds site directed PCR mutagenesis on pCZGY3010, first using primers YJ12463 and YJ12464, then primers YJ12465 and YJ12466. To generate *Pref-1*::*eif-3.I*(+) (pCZGY3508), we amplified *eif-3.I* cDNA from N2 cDNA libraries using primers YJ12453 and YJ12454, and used Gibson Assembly to clone into the pCZGY66 backbone containing *Prgef-1*. We then performed site-directed mutagenesis on pCZGY3508 using primers YJ12457 and YJ12458 to generate the *Prgef-1*::*eif-3.I*(*Q252R*) construct (pCZGY3509).

We generated the GFP::EIF-3.G clones pCZGY3018 and pCZGY3019 via Gibson assembly^®^, using *eif-3.G*(+) or *eif-3.G*(*C130Y*) cDNA amplified using primers YJ12604 and YJ12605, and the GFP-coding DNA amplified using primers YJ12602 and YJ12603.

To generate *Punc-17β*::EIF-3.G::3xFLAG::SL2::GFP constructs (pCZGY3538 for WT, pCZGY3539 for C130Y, and pCZGY3540 for ΔRRM) used in seCLIP experiments, *Punc-17β* promoter was amplified from pCZGY1091 using primers YJ12164 and YJ12418, each *eif-3.G* cDNA (wild type, C130Y, or ΔRRM) was amplified with an N-terminal 3xFLAG sequence from subclones using the primers YJ12419 and YJ12420, SL2 trans-splice sequence was amplified from N2 genomic DNA using primers YJ12421 and YJ12422, and GFP was amplified from pCZGY3018 using primers YJ12423 and YJ12424. These fragments were then Gibson Assembled into the pCZGY2729 backbone, which facilitates CRISPR/Cas9 single copy insertion on chromosome IV (Andrusiak et al. 2019).

All *ncs-2* transgenes were similarly cloned using primers for Gibson assembly into pCZGY2727. To generate the *Pncs-2*::5′UTR mutant::*ncs-2* cDNA construct (pCZGY3526), we amplified *Pncs-2* from N2 gDNA using primers YJ12554 and YJ12555. A fragment containing SL1 trans-spliced *eif-3.G* 5′UTR incorporated in the forward primer, *ncs-2* cDNA, GFP, and the *ncs-2* 3′UTR was amplified from CZ22459 gDNA using primers YJ12556 and YJ12557. The *Pncs-2*::GFP(+) construct (pCZGY3533) was cloned by amplifying *Pncs-2* through the first four codons of *ncs-2* CDS from N2 gDNA using primers YJ12554 and YJ12579, and GFP and the *ncs-2* 3′UTR from CZ22459 gDNA using YJ12580 and YJ12557. The *Pncs-2*::5′UTR mutant::GFP construct (pCZGY3534) was cloned by amplifying *Pncs-2* from N2 gDNA using primers YJ12554 and YJ12555, and the *eif-3.G* 5′UTR, the first four codons of the *ncs-2* CDS, GFP, and the *ncs-2* 3′UTR from CZ22459 gDNA using primers YJ12581 and YJ12557.

### Fluorescence microscopy and GFP intensity quantification

L4 or young adult animals were immobilized in 1mM levamisole in M9 and mounted on microscope slides with 2% agar. All images were collected on a Zeiss LSM800 confocal microscope, unless specified, with identical image acquisition settings: 1.25 µm pixel size with 0.76 µs pixel time, 50 µm pinhole, with genotype-blinding to observer when possible. The positions of VA10, VB11, and DB7 cholinergic motor neurons were identified using *juEx2045*(*Pacr-2-mCherry*), based on their stereotypical patterning in the posterior ventral nerve cord. These neurons were chosen for quantification because they were consistently visible in single focal plane images. All quantification of GFP intensity in these neurons was performed using the Integrated Density function in ImageJ (Schindelin et al. 2012). We acquired the mean integrated density from the VA10, VB11, and DB7 cell bodies, subtracted background intensity from an equivalent area, and the resulting values were then normalized to the mean area of the cell bodies of the same animal. We similarly quantified fluorescence intensities in the ventral nerve cord of animals expressing GFP-tagged full-length *ncs-2* cDNA, except integrated densities were obtained from one ROI per image (red boxes in Fig. 7B and 7C). All data was normalized to the mean fluorescence intensity of the transgene in the wildtype background. All statistical analysis was performed with GraphPad Prism6 software.

Axon commissures, observed as fluorescent structures extending from the ventrally located neuron cell body to the dorsal body wall, shown in Supplemental Fig. S2A were visualized with *juIs14*[*Pacr-2*::GFP] and manually quantified. Imaging shown in Supplemental Fig. S2B was performed using a Zeiss Axioplan 2 microscope installed with Chroma HQ filters and a 63x objective lens. Synaptic puncta labeled by *nuIs94*[SNB-1::GFP], were manually quantified in the region anterior to the ventral nerve chord between VD6 and VD7.

### Polysome Profiling

We prepared *C. elegans* lysates and sucrose gradients using the protocol described in (Ding and Grosshans 2009). To synchronize animals, gravitated adults were treated with 20% Alkaline Hypochlorite Solution and embryos were plated on four 30 cm NGM plates seeded with OP50, and grown to the L4 stage at 20°C. Approximately 200 µl packed L4 *C. elegans* were harvested by centrifugation in M9 media at 1,500 RPM, washed three times in ice-cold M9 media supplemented with 1mM cycloheximide, then once more in lysis buffer base solution (140 mM KCl, 20 mM Tris-HCl (pH 8.5), 1.5 mM MgCl_2_, 0.5% NP-40, 1 mM DTT, 1 mM cycloheximide) followed by snap freezing in liquid nitrogen. The frozen pellets were resuspended in 450 µl lysis buffer (140 mM KCl, 20 mM Tris-HCl (pH 8.5), 1.5 mM MgCl2, 0.5% NP-40, 2% PTE, 1% sodium deoxycholate, 1 mM DTT, 1 mM cycloheximide, 0.4 units/µl RNAsin) and crushed to a fine powder with a mortar and pestle pre-cooled with liquid nitrogen. Protein lysate concentrations were then determined using a Bradford assay (Bio-Rad, Hercules, CA). 15-60% sucrose gradients were prepared in 89 mm polypropylene centrifuge tubes (Beckman Coulter) using standard settings on a Foxy Jr. density gradient fractionation system (Teledyne ISCO, Lincoln, NE) and lysate volumes corresponding to equal protein amounts between samples were loaded on top of the gradients. Loaded gradients were then spun in an Optima L-80 ultracentrifuge (Beckman Coulter) at 36,000 rpm at 4°C for 3 hours. Fractions were then collected and RNA absorbance was continuously acquired using a UA-6 detector (Teledyne ISCO, Lincoln, NE) with a 70% sucrose chase solution. We calculated the area under the curve (AUC) for monosome (80S) and polysome absorbance traces using the Simpson’s rule method in SciPy (Virtanen et al. 2020) and used the AUC values to calculate the polysome to monosome ratios.

### seCLIP library preparation and sequencing

We performed single-end enhanced CrossLink and ImmunoPrecipitation (seCLIP) experiments according to the published protocol in (Van Nostrand et al. 2017), with the following adjustments to ensure efficient immunoprecipitation yield from *C. elegans* lysates. Mixed stage animals were grown on ∼12 NGM plates (30 cm) and washed twice with M9, spinning at 1,500 rpm between washes. Animals were then resuspended in 5 ml M9 media and rocked on a rotator for 10 minutes to remove gut bacteria, followed by one more wash with M9 at 1,500 rpm. The animals were spread on one NGM plate (30 cm) and then UV-crosslinked with a Spectrolinker™ XL-1000 (Spectronics, New Cassel, NY) using energy setting 3 kJ/m^2^ according to (Broughton and Pasquinelli 2013). Afterwards, animals were resuspended in 4 ml lysis buffer [150 mM NaCl, 1 M HEPES, 100 mM DTT, 6.25 µl RNAsin (Promega) per 10 ml, 10% glycerol, 10% Triton X-100, 1 protease inhibitor tablet per 10 ml] and split into two tubes for each replicate. The resuspension was disrupted on an XL-2000 Sonicator (QSonica, Newtown, CT) with 7 pulses (powersetting= 11, 10 seconds each, 50 seconds on ice in between) and immediately spun at 4,750 RPM for 5 minutes at 4°C. All subsequent steps, beginning with RNAse A treatment of the supernatant, was performed according to the seCLIP protocol (Van Nostrand et al. 2017), except that high salt and low salt wash buffers were replaced with a single buffer (2M NaCl, 1M HEPES, 30% glycerol, 1% Triton X-100, 1 protease inhibitor tablet per 10 mL) optimized for anti-FLAG RNA IP from *C. elegans* lysates (Blazie et al. 2015). Immunoprecipitation was performed with anti-FLAG beads (Sigma, St. Louis, MO).

cDNA libraries were prepared from both the immunoprecipitated mRNA (CLIP) as well as the sample before immunoprecipitation (INPUT), such that crosslink sites can be defined by read enrichment in the CLIP sample over input as described (Van Nostrand et al. 2017). seCLIP libraries were validated using the D1000 high sensitivity screen tape system (Agilent, La Jolla, CA) and quantified using a Qubit instrument (Thermo Fisher, San Diego, CA) before pooling and sequencing on HiSeq4000 (Illumina, San Diego, CA) at the IGM Genomics Center, University of California San Diego.

### seCLIP sequence mapping

After demultiplexing barcoded reads, we used the CLIPPER software pipeline (Lovci et al. 2013) to trim barcodes, remove PCR duplicate reads, filter reads mapping to repetitive elements, and map the remaining reads to the *C. elegans* reference genome (ce10). The total number of uniquely mapped reads obtained after filtering is in Supplemental Table S4. A large proportion of reads obtained from the ΔRRM and IgG samples mapped to repetitive elements and were discarded, explaining the smaller number of uniquely mapped reads in these samples. In seCLIP, RNA binding sites are defined as read clusters enriched in the crosslink immunoprecipitated sample (CLIP) over the input control (INPUT) (Van Nostrand et al. 2017), which are comprehensively identified across each dataset using CLIPPER. Read clusters were reproducibly identified from independent biological replicates of seCLIP, except in the ΔRRM control reflecting background, supporting the specificity of our data (Supplemental Fig. S4).

### EIF-3.G footprint identification from mapped seCLIP reads

We defined EIF-3.G footprints as seCLIP read clusters appearing in both replicates with 20 reads and 1.5 fold-change enrichment over the INPUT control in at least one replicate. Footprints matching these criteria in the IgG (no transgene) and the EIF-3.G(ΔRRM) control samples were considered background and subtracted from the EIF-3.G(WT) and EIF-3.G(C130Y) datasets (Supplemental Tables S5 and S6). We annotated footprints to their gene features (eg. 5′UTR, CDS) using custom scripts that overlap read clusters with the *C. elegans* genome annotation WS235.

### Gene Ontology and KEGG pathway analysis

GO analysis was performed using 225 EIF-3.G target genes as input to the biological process annotation set within the Gene Ontology Resource tool (Ashburner et al. 2000). A total 211 gene names were recognized by the database and GO term enrichment was defined using a threshold of P< 0.05. Pathway analysis of EIF-3.G target genes was performed using the Kyoto Encyclopedia of Genes and Genomes (KEGG) annotation tool within the DAVID bioinformatics resource (Jiao et al. 2012) using default settings.

### Analysis of activity-dependent expression changes among EIF-3.G target mRNAs in cholinergic neurons

We studied activity-dependent transcript expression changes among the EIF-3.G target genes (n= 225) by re-analyzing the cholinergic neuron-specific transcriptomes reported in (McCulloch et al. 2020) using the Galaxy platform (Afgan et al. 2018). We downloaded raw FASTA reads from transcriptome sequencing of wild type and *acr-2*(*gf*) animals (n= 2 replicates each; accession #’s SRR10320705, SRR10320706, SRR10320707, SRR10320707) and mapped them to the *C. elegans* reference genome (ce10) using BWA (Li and Durbin 2009). Differential expression among the EIF-3.G target genes was quantified using Feature Counts (Liao et al. 2014) and DeSeq2 (Love et al. 2014).

### *in silico* analysis of 5′UTR sequence features, secondary structure, and conservation

We downloaded all *C. elegans* transcript 5’UTRs (WS271) from Parasite Biomart (Howe et al. 2016), and calculated 5′UTR lengths as the sequence between the 5’ distal end and the start codon of each transcript. To have meaningful length calculation, we only considered 5′UTRs annotated with at least 10nt and restricted our analysis to the longest 5′UTR isoform for each gene to avoid considering multiple transcripts of the same gene. By these criteria we identified 5′UTRs for 10,962 WS271 protein coding transcripts and 179 transcripts with EIF-3.G footprints.

For the analysis shown in Supplemental Fig. S5D and S5E, the genomic coordinates of human gene 5′UTRs were downloaded from Ensembl and used to obtain 5′UTR sequences from the human genome reference sequence (hg38). eIF3g footprints from HEK293 cells were previously described (Lee et al. 2015). We defined our analysis of 5′UTR sequences using the same criteria described for *C. elegans* and the data show the comparison between 5′UTRs of 255 genes with human eIF3g footprints and 19,914 total genes in the human genome annotation (hg38).

To calculate GC-enrichment, we used BEDTools (Quinlan and Hall 2010) to generate a FASTA of 5′UTR sequences from their genomic coordinates and used a custom Python script to calculate the total %GC in their sequences (Figs. 4F) as well as %GC within 10nt bins incremented from the start codon ATG for the analysis shown in Supplemental Fig. S5C.

To predict secondary structures of the 5′ ends of *hlh-30d*, *ncs-2*, and *eif-3.G* mRNAs, we used the RNAfold Web Server (Gruber et al. 2008) with default settings. To better understand the contribution of gene-specific 5’UTR sequences, we excluded the SL1 sequences of *ncs-2* and *eif-3.G* from folding predictions. The free energies (ΔG) for each sequence reported in our results were derived from the reported thermodynamic ensemble. Data showing conservation of *eif-3.G* and *ncs-2* 5′UTR sequences compared with 135 nematode species (phyloP135way scores) was obtained from the UCSC Genome Browser with the genomic position along the sequence of each 5′UTR as input.

## Data accessibility

Raw and analyzed seCLIP datasets from this study have been uploaded to the Gene Expression Omnibus (GEO) under accession number GSE152704.

## Acknowledgements

We thank Brian Yee, Eric Van Nostrand, Gabriel Pratt, and Gene Yeo for advice on adapting the seCLIP protocol to *C. elegans* and technical support with seCLIP data analysis; Timothy Shaw and Jens Lykke-Andersen for sharing equipment and invaluable assistance with polysome profile analysis; Yan Zhao and Ann Zhou for assistance in constructing expression clones and strains; Malena Hansen, Josh Kaplan, and Keming Zhou for transgenes, Kenneth Miller for P*unc-17β* plasmid. Some strains used in this study were provided by the Caenorhabditis Genetics Center, which is funded by NIH Office of Research Infrastructure Programs (P40 OD010440). We acknowledge WormBase and UCSC Genome databases for genomic information resource. We thank Andrew D. Chisholm and members of our laboratory for critical reading of the manuscript. S. B. was a trainee on the UCSD T32 training grant (NS007220). This work is funded by NIH grant (NS R37 035546 to Y. J.)

## Author Contributions

SMB, STK, YJ conceived the project and designed experiments. SMB and STK performed experiments and data analyses. KM contributed expression data. SMB and YJ wrote manuscript, STK contributing to manuscript editing. All authors read the manuscript.

---

**Supplemental Fig. S1:**
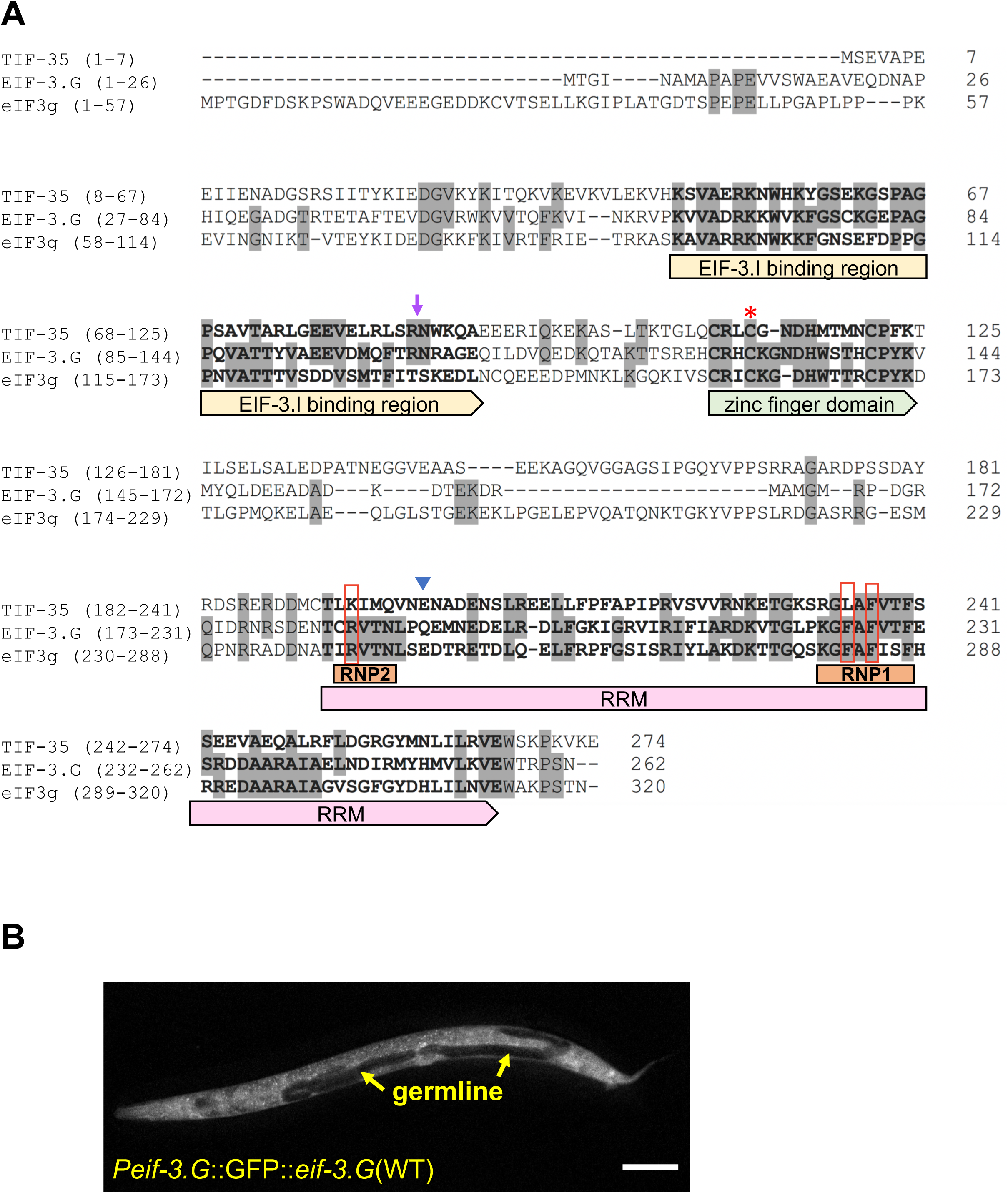
EIF-3.G is highly conserved and expressed ubiquitously. **A)** Clustal-W sequence alignment of *C. elegans* EIF-3.G (NP_001263666.1) with the *S. cerevisiae* (TIF-35; 32% identity; NP_010717.1) and human (eIF3g, 35% identity; AAC78728.1) orthologs. Residues identical to *C. elegans* EIF-3.G are shaded gray. The EIF-3.I binding region, zinc finger, RRM and RNP motifs are indicated below the corresponding sequences. The frame-shift caused by the *ju1327* deletion adds 85aa’s starting from the position marked by the purple arrow. The C130Y mutation (red asterisk), R-F-F residues changed to alanine in our RFF/AAA transgene construct (red boxes), and location of the Q191* mutation used to generate EIF-3.G(ΔRRM) (blue arrow head) are shown. **B)** Representative single plane confocal image of an L4 stage animal expressing GFP::EIF-3.G(WT) under the *Peif-3.G* promoter. GFP is visualized throughout *C. elegans* tissues and excluded from the gonads due to germline transgene silencing. Scale bar = 100 µm.

**Supplemental Fig. S2:**
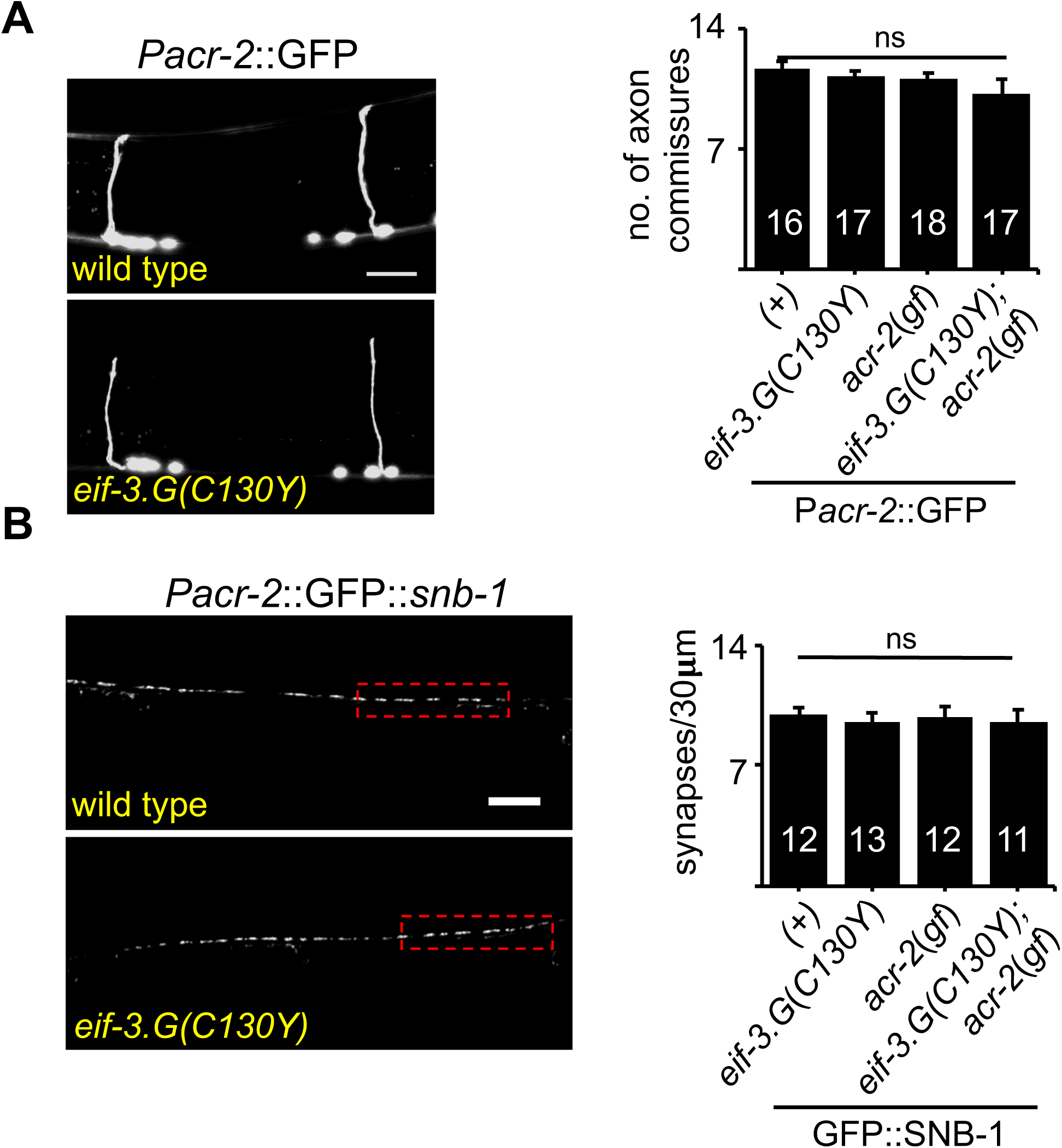
Motor neuron development is normal in *eif-3.G*(*C130Y*) animals. **A)** Representative confocal z-stack projection images of axonal commissures projecting from motor neurons of wild type or *eif-3.G*(*C130Y*) single mutant L4 animals (head to the right). The quantification of axon commissure numbers in strains is shown on the right. **B)** Single plane confocal images of L4 animals (head to the left) expressing GFP::SNB-1, which appear as puncta along the ventral nerve cord. Puncta quantification, shown at right, was performed in a 30 µm region anterior of the vulva (red box) in each animal. For **A)** and **B)**, the number of animals used in quantification data is indicated in each bar and error bars represent ±SEM. Scales bar = 10 µm.

**Supplemental Fig. S3:**
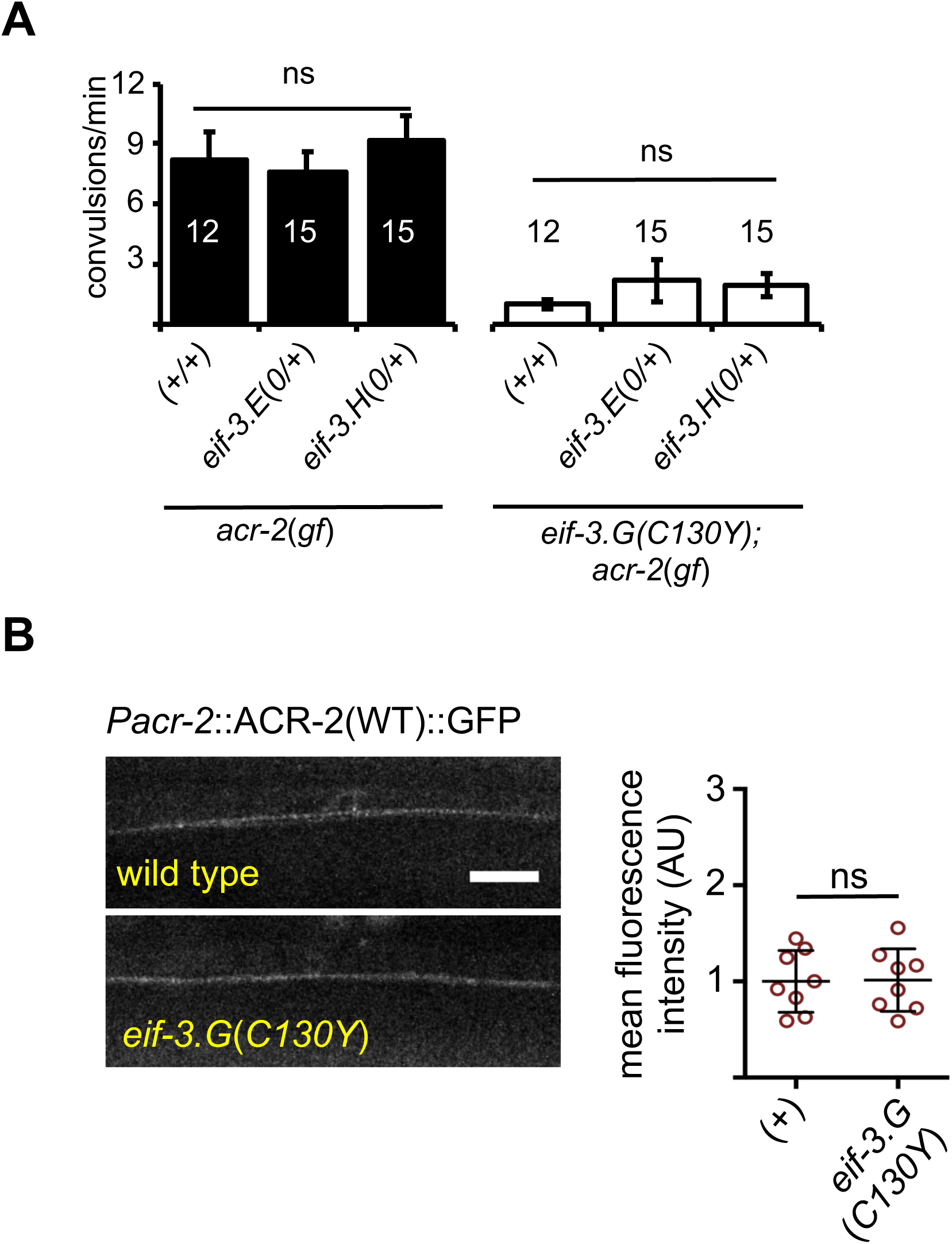
EIF-3.G(C130Y) modulation of convulsion behavior does not involve reduced EIF-3 complex dosage or ACR-2 expression. **A)** Quantification of convulsion frequencies in animals of the indicated genotypes. Error bars are ± SEM and n= 15 per sample. (ns) not significant by one-way ANOVA with Bonferroni’s post-hoc test. **B)** Representative images of L4 animals expressing ACR-2(WT)::GFP in the ventral nerve chord region anterior to the vulva. Quantification of fluorescence in animals (n= 8) is shown on the right. Scale bar = 10 µm. (ns) not significant by two-tailed t-test.

**Supplemental Fig. S4:**
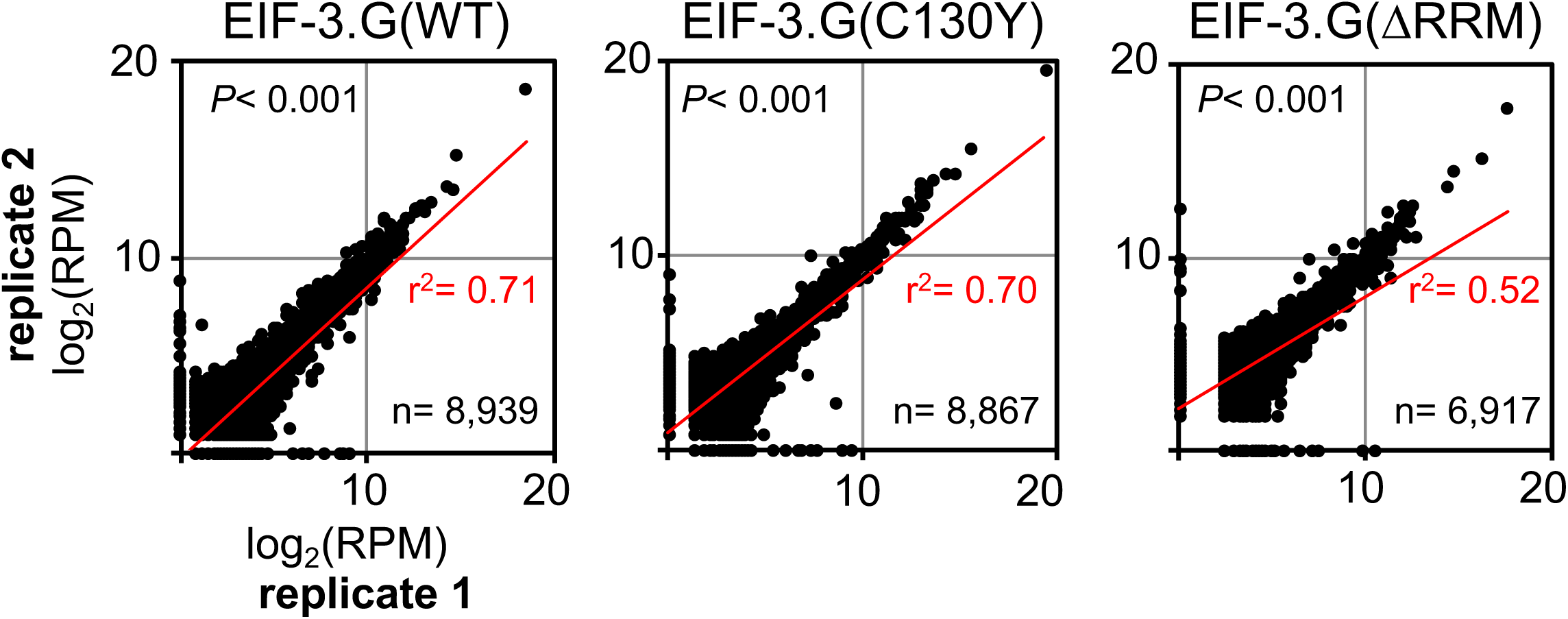
Replicate seCLIP experiments produce similar results. Scatter plots comparing the cumulative RPM (reads per million mapped reads) mapped in genes detected between seCLIP replicates. Each dot represents a unique gene and RPM values are the total reads for all clusters mapped within the respective gene. P-values are derived from a two-tailed Pearson’s correlation (r^2^). The number of genes per dataset (n) and linear fit (red line) are shown.

**Supplemental Fig. S5:**
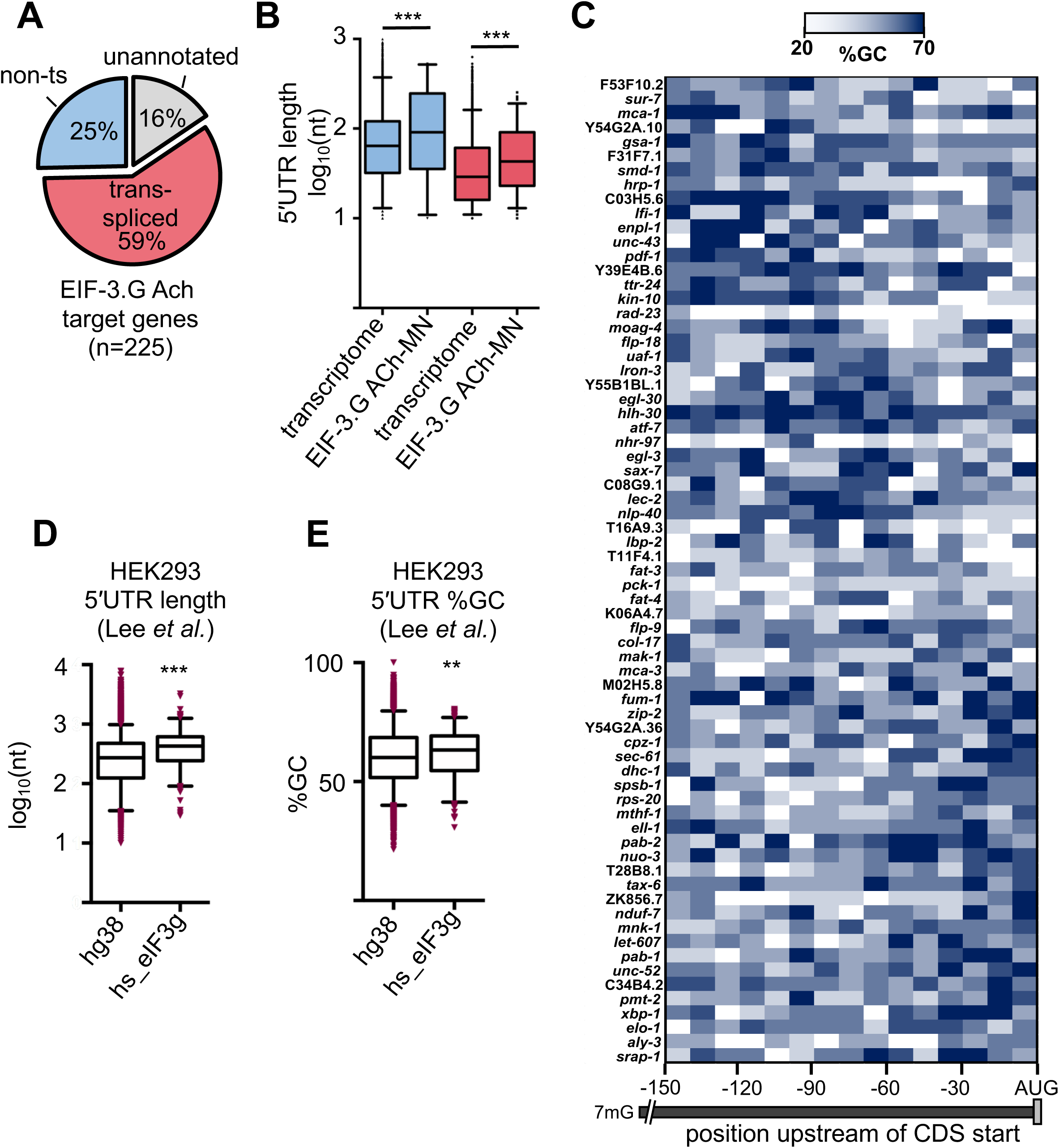
EIF-3.G associates with long and GC-rich 5′UTRs. **A)** Pie charts show the proportion of trans-splicing (ts) among genes with 5′UTR proximal EIF-3.G footprints according to Allen *et al*. Unannotated genes were absent from the Allen *et al*. dataset. **B)** Box plots compare the lengths of non-trans-spliced (blue) or trans-spliced (red) 5′UTR sequences of EIF-3.G target mRNAs (non-trans-spliced n= 57, trans-spliced n= 133) to all 5′UTRs in the *C. elegans* transcriptome (WS271; non-trans-spliced n= 5,970, trans-spliced n= 6,674). Boxes are 10-90 percentile with outliers in red. **C)** Heat map shows 5′UTR %GC along the first 150 nts from the start codon in 10 nt bins of EIF-3.G target genes. Only genes with a 5′UTR of at least 150 nts (n= 49) are shown. Genes are clustered top to bottom by increasing positional GC-density near the start codon. **D) and E)** Box plots comparing **D)** length or **E)** GC-content of 5′UTR sequences of HEK293 eIF3g target mRNAs (n= 255; Lee *et al.,* 2016) versus all human transcriptome 5′UTRs (hg38; n= 19,914). Boxes are 5-95 percentile with outliers in red. For **B**, **D**, and **E)** Statistics: (***) P< 0.001, (**) P< 0.01, (ns) not significant by two-tailed Mann-Whitney test.

**Supplemental Fig. S6:**
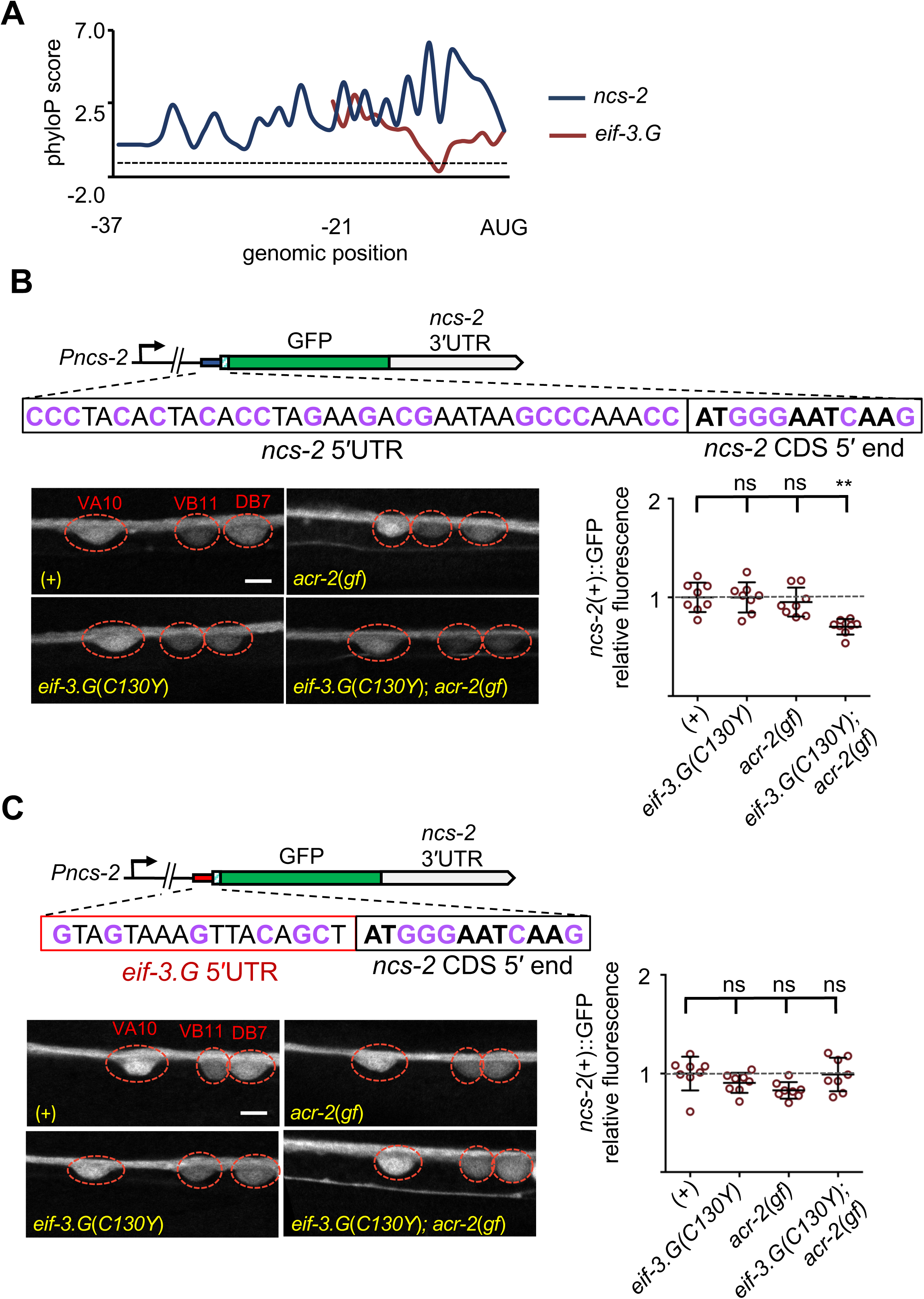
EIF-3.G(C130Y) reduces NCS-2 expression in the ACh-MNs of *acr-2*(*gf*) animals dependent on its conserved 5′UTR. **A)** Plot shows conservation of nucleotides (by phyloP135way scores) in the 5′UTR sequences of *ncs-2* (blue) and *eif-3.G* (red), with higher scores indicating greater conservation. Genomic position (x-axis) is relative to the start codon (AUG). The horizontal dashed line marks PhyloP score= 0. **B)** Top: Schematic of the *Pncs-2*::GFP translation reporter containing the endogenous GC-rich (purple) *ncs-2* 5′UTR and the first 4aa of NCS-2 followed by GFP and driven by the *Pncs-2* promoter. *Bottom:* Representative single-plane confocal images of *Pncs-2*::GFP expression in the VA10, VB11, DB7 soma in animals of the indicated genotypes. GFP intensity quantification are shown to the right (n= 8). **C)** Top: The *Pncs-2*(5′UTR mutant)::GFP translation reporter contains the 5′UTR of *eif-3.G* (red, GC content is purple), followed by the *ncs-2* CDS 5’ end. *Bottom:* Representative single-plane confocal images of *Pncs-2*(5′UTR mutant)::GFP in VA10, VB11, DB7 soma in the indicated genetic backgrounds. GFP intensity quantification are shown to the right (n= 8). For GFP quantification in panels **B)** and **C)**, data points are normalized to the mean intensity of the respective reporter in the wild type background. Red dashes indicate positions of labeled ACh-MN soma. Scale bar = 15 µm. Statistics: (***) P< 0.001, (ns) not significant by one-way Anova with Bonferroni’s post hoc test.

**Supplemental Video S1: N2 [Wild type] *C. elegans* movement on solid nematode growth media.**

**Supplemental Video S2: MT6241 [*acr-2*(*gf*)] *C. elegans* movement on solid nematode growth media.**

**Supplemental Video S3: CZ21759 [*eif-3.G*(*C130Y*); *acr-2*(*gf*)] *C. elegans* movement on solid nematode growth media.**

**Supplemental Video S4: CZ22197 [*eif-3.G*(*C130Y*)] *C. elegans* movement on solid nematode growth media.**

**Supplemental Table S1:**
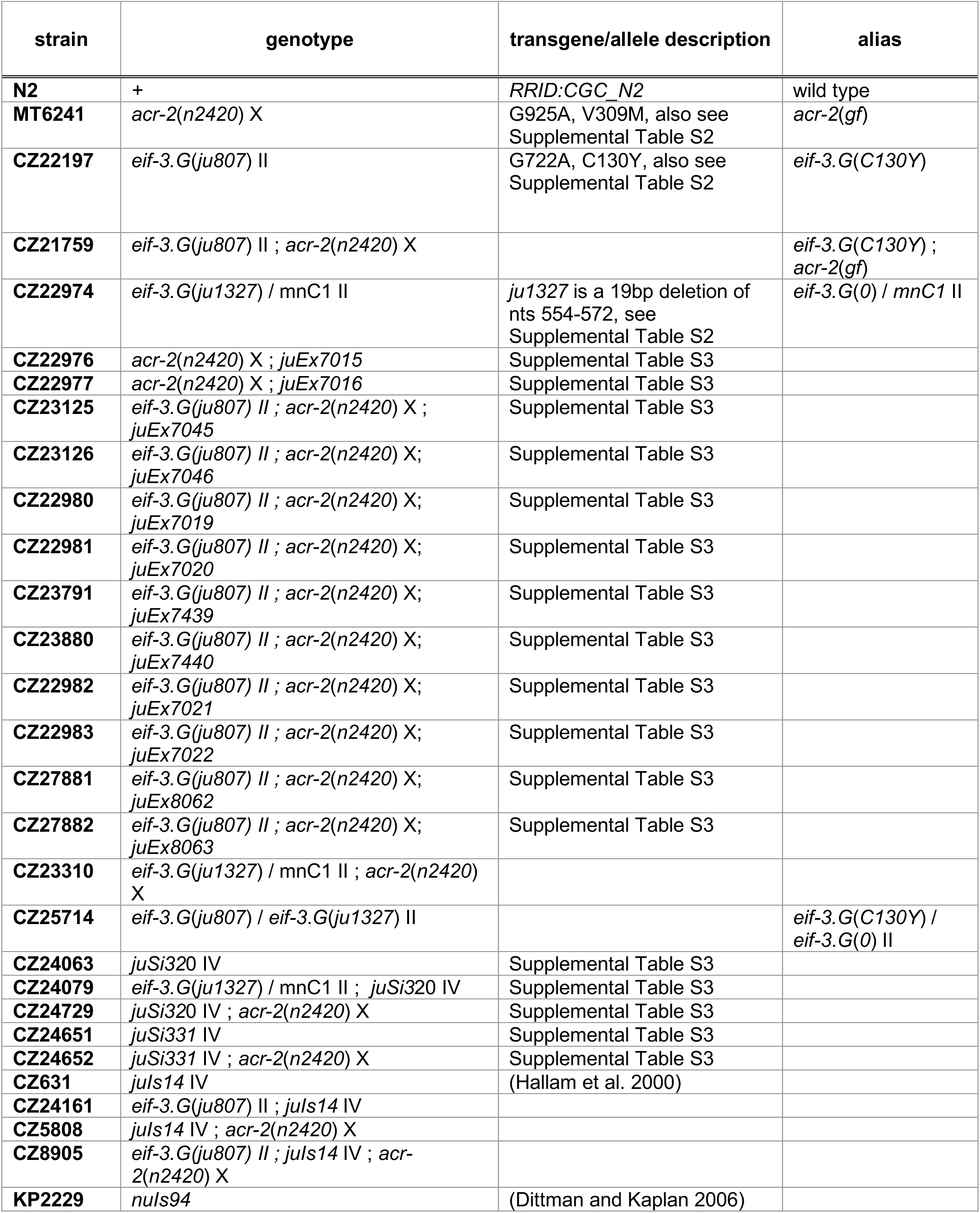
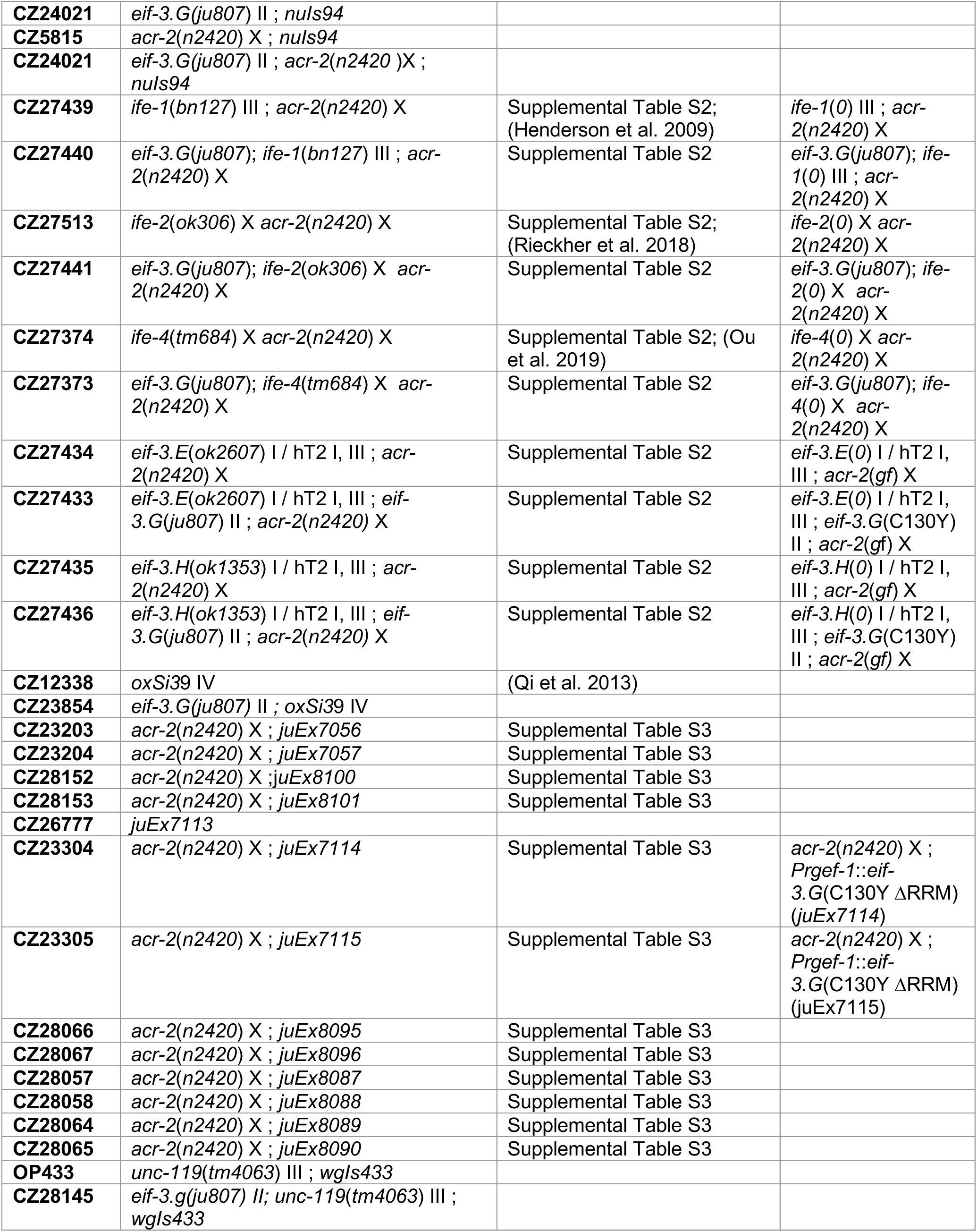
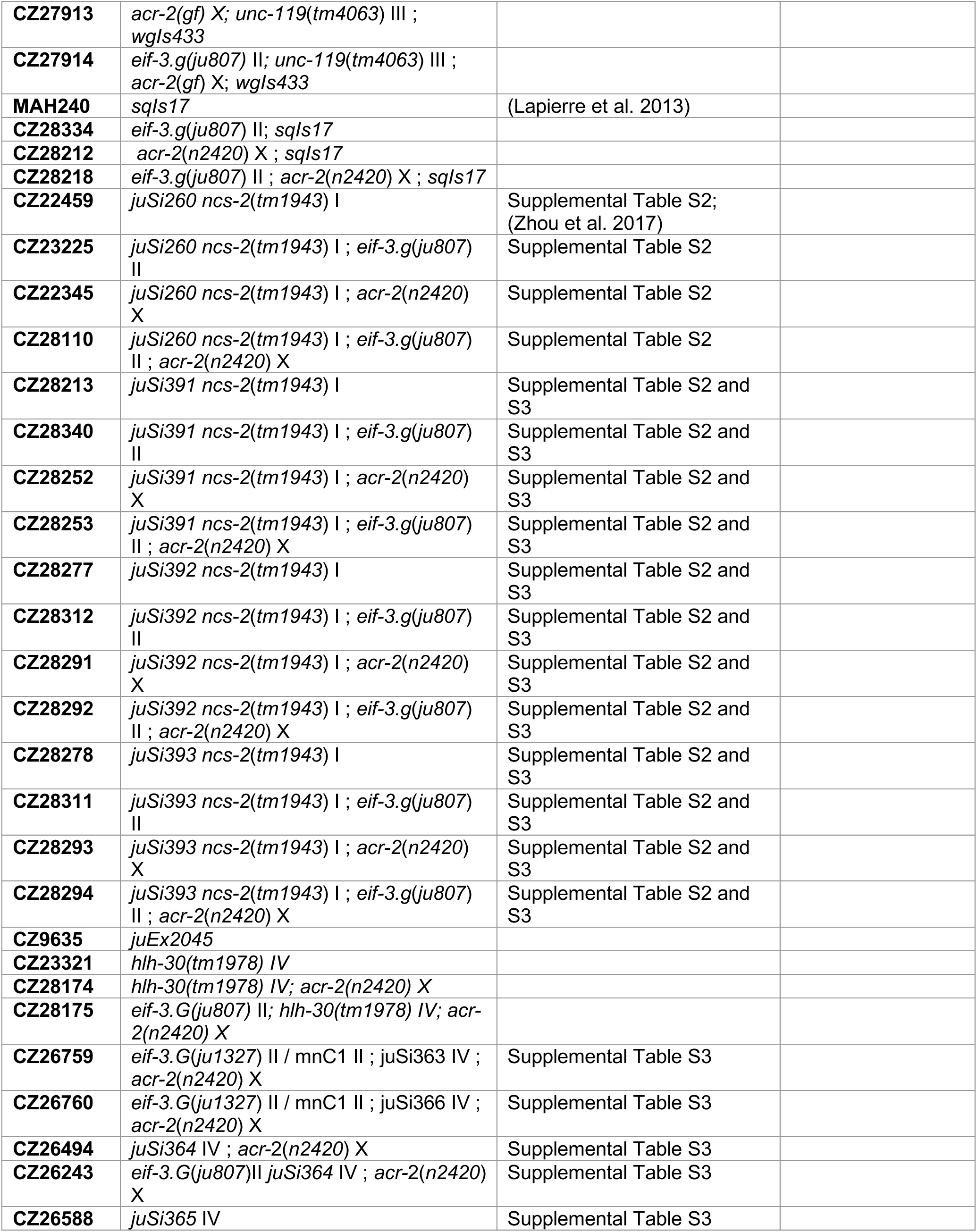
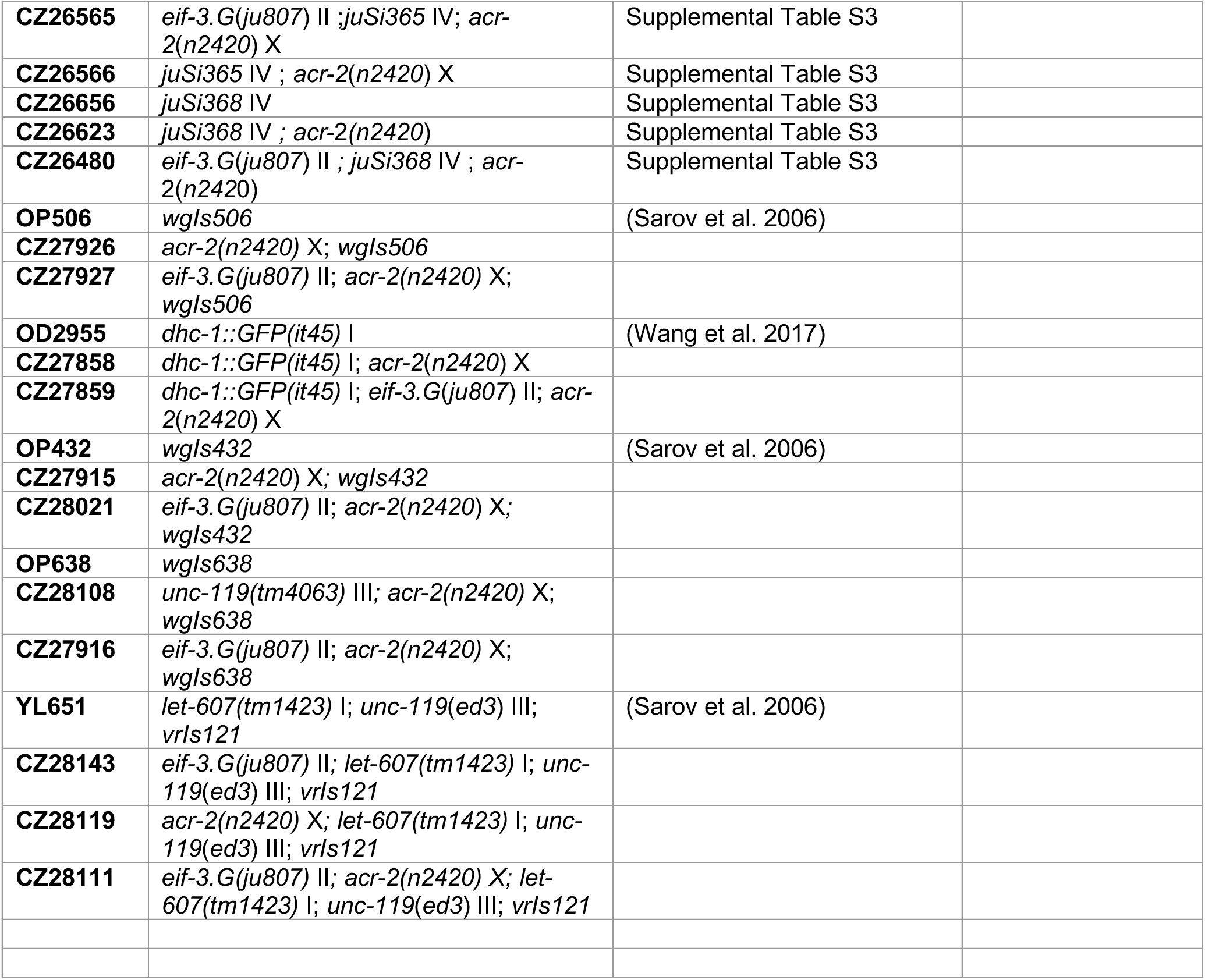
Strains used in this study.

**Supplemental Table S2:**
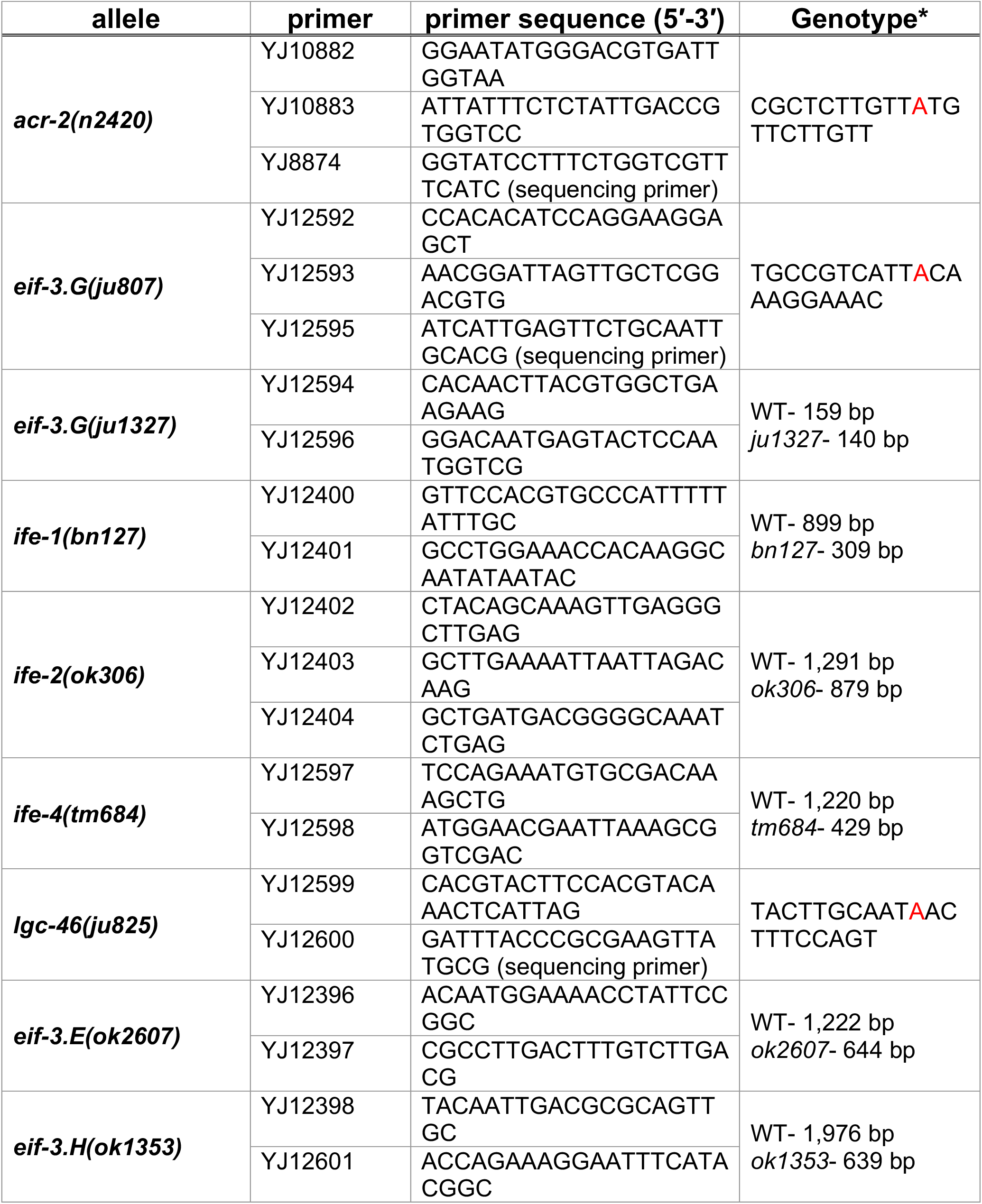
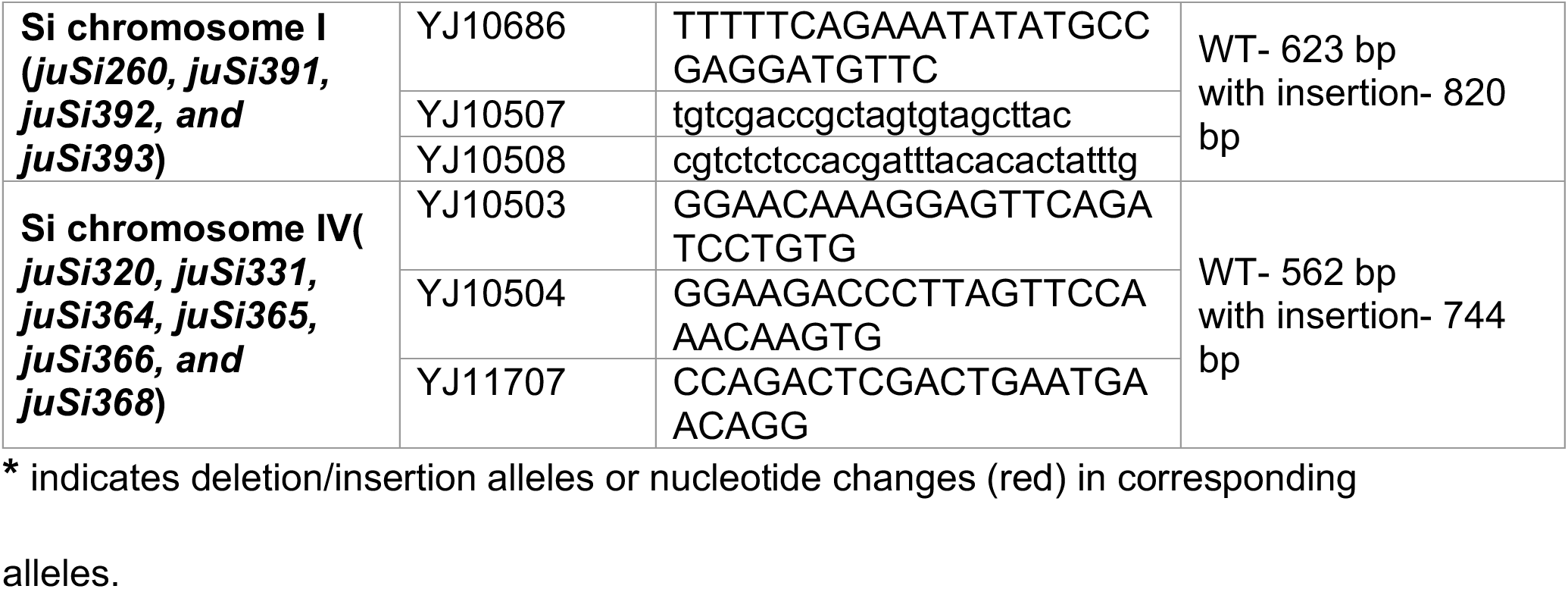
Genotyping primers used in this study.

**Supplemental Table S3:**
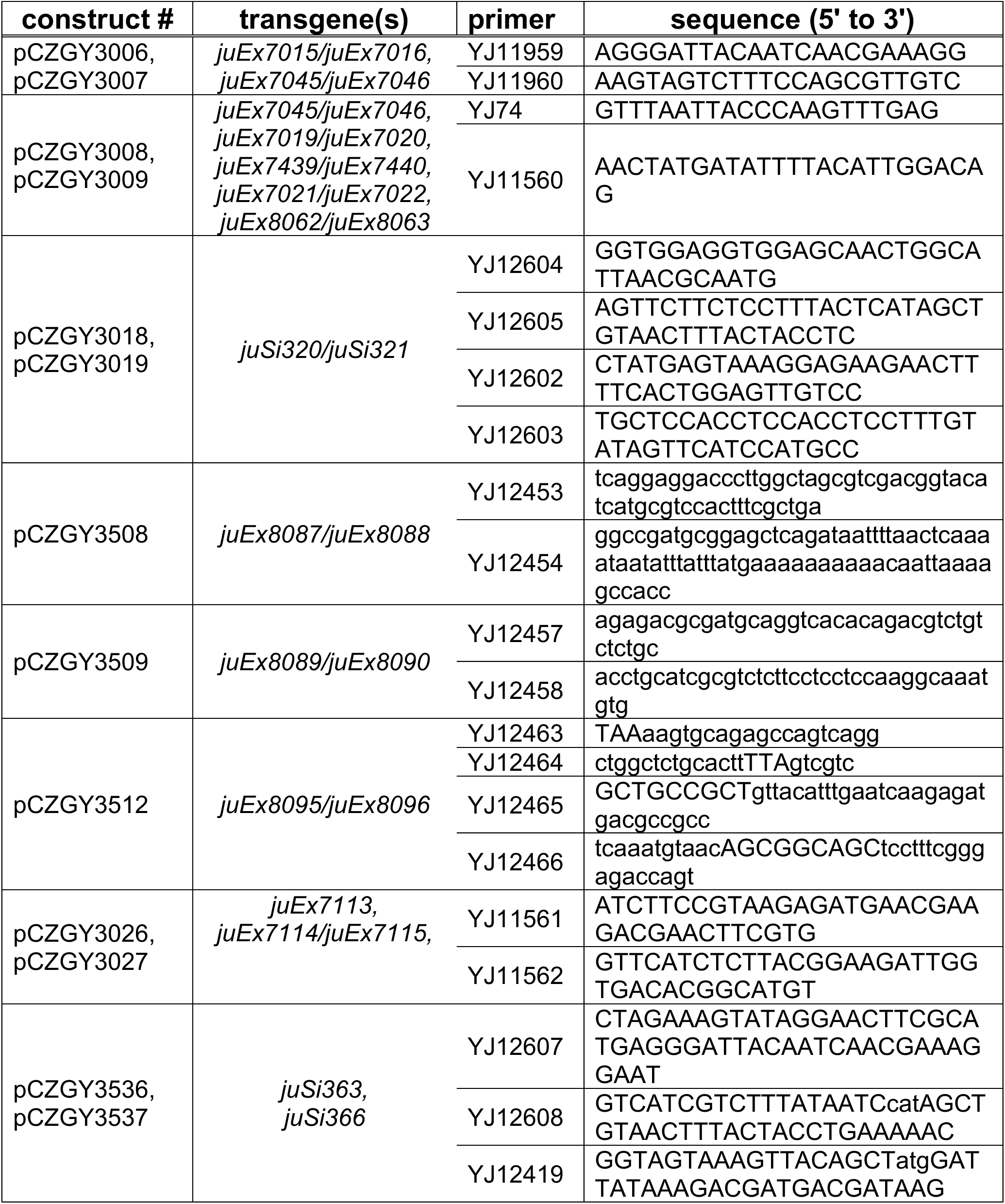
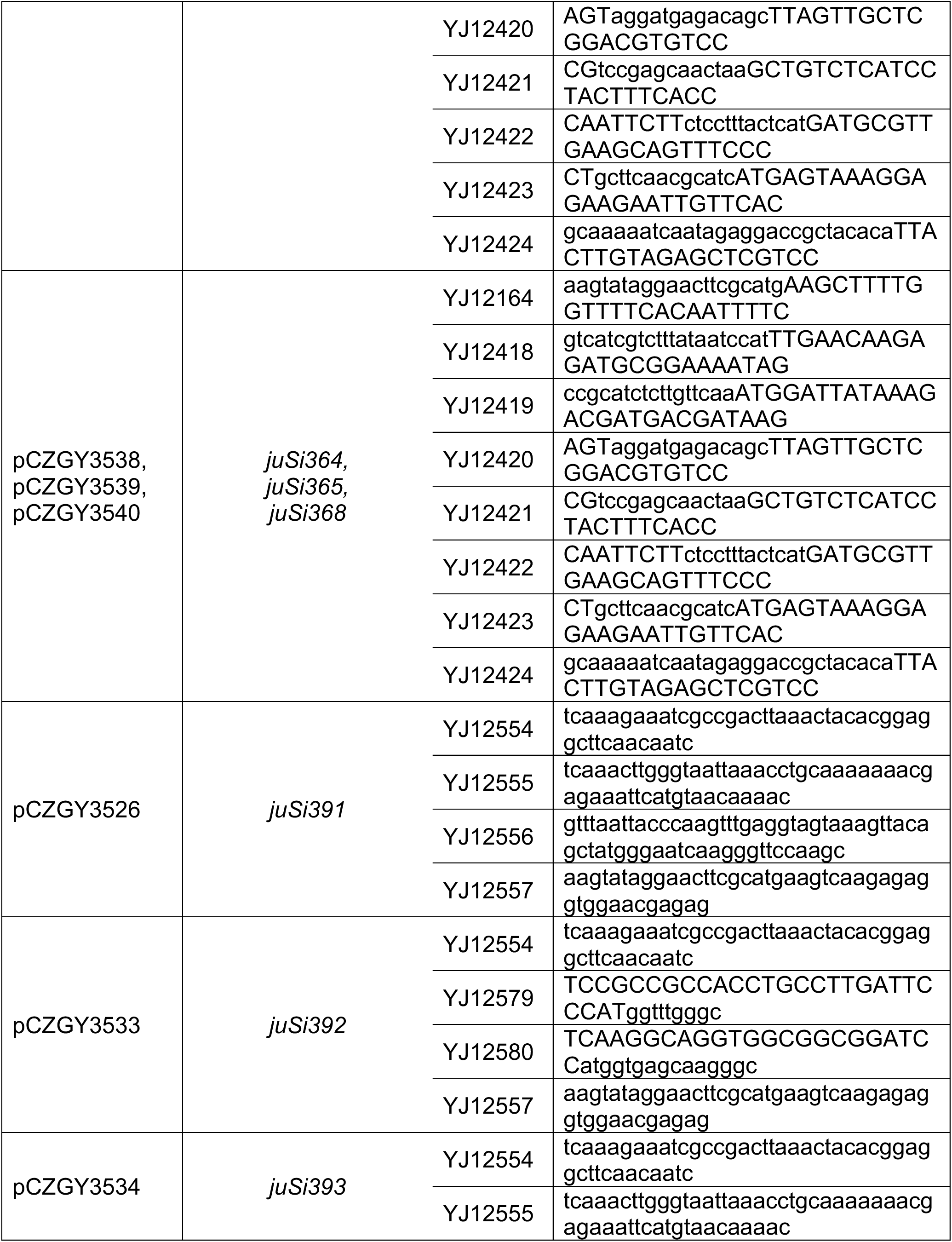
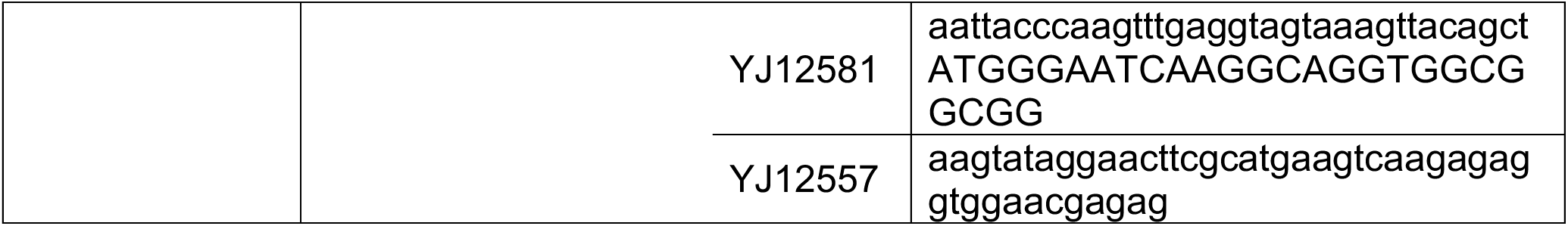
Constructs and related primers used in this study.

**Supplemental Table S4:**
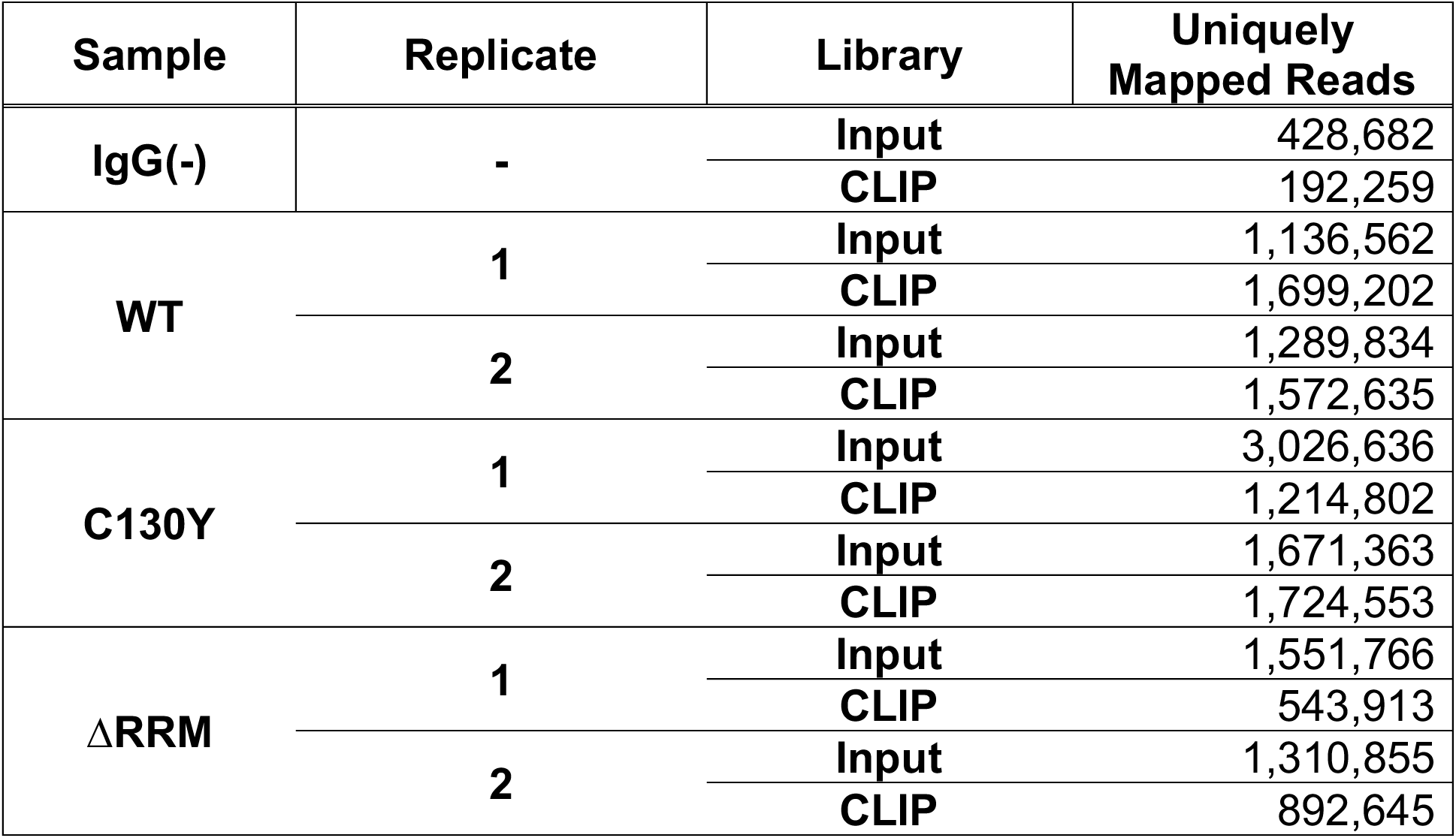
Number of mapped reads in seCLIP replicate datasets obtained after sequencing and CLIPPER filtering.

| Sample | Replicate | Library | Uniquely Mapped Reads |
| --- | --- | --- | --- |
| IgG(-) | - | Input | 428,682 |
|  |  | CLIP | 192,259 |
| WT | 1 | Input | 1,136,562 |
|  |  | CLIP | 1,699,202 |
|  | 2 | Input | 1,289,834 |
|  |  | CLIP | 1,572,635 |
| C130Y | 1 | Input | 3,026,636 |
|  |  | CLIP | 1,214,802 |
|  | 2 | Input | 1,671,363 |
|  |  | CLIP | 1,724,553 |
| ΔRRM | 1 | Input | 1,551,766 |
|  |  | CLIP | 543,913 |
|  | 2 | Input | 1,310,855 |
|  |  | CLIP | 892,645 |

**Supplemental Table S5:**
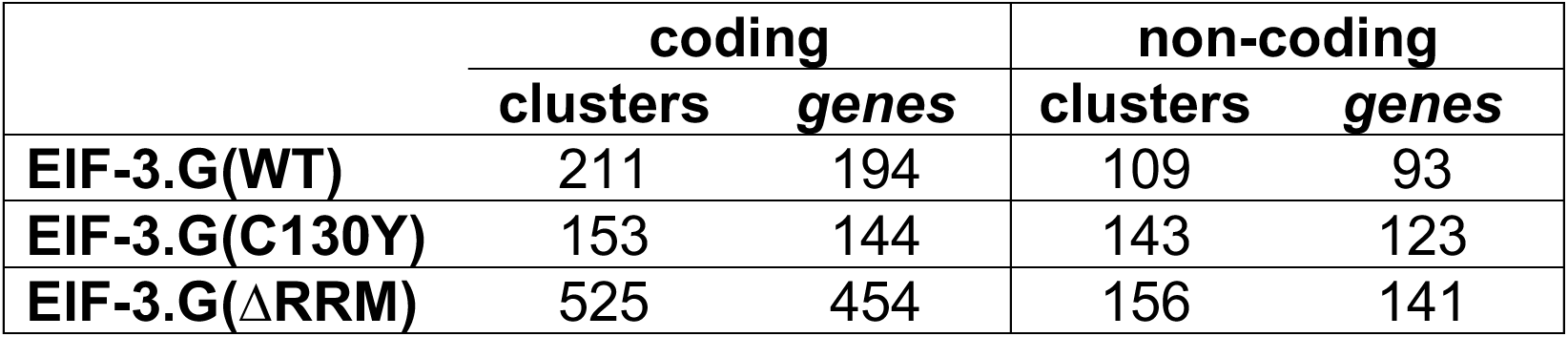
Number of read clusters detected in each dataset after subtraction of IgG control background.

**Supplemental Table S6:**
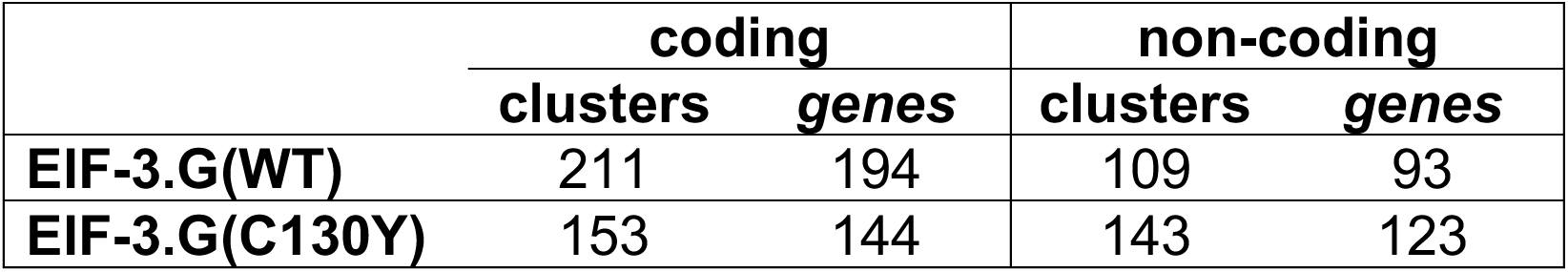
Number of EIF-3.G footprints detected in each dataset after subtraction of background from both IgG and ΔRRM controls.

